# Galectin-9 drives thrombo-inflammatory circuits initiating venous thrombosis

**DOI:** 10.64898/2026.11.05.757779

**Authors:** Akhil A. Konkoth, Adel Abo Mansour, Katharine Whitworth, Ivan Budnik, Daniel Steiner, Cihan Ay, Danielle Lezama, Helen M. McGettrick, Dean Kavanagh, Francesco Maione, Juma El-Awaisi, G. Ed Rainger, Julie Rayes, Alexander Brill, Asif J. Iqbal

**Affiliations:** Department of Cardiovascular Sciences, College of Medicine and Dental Sciences, School of Medical Sciences, University of Birmingham, Birmingham B15 2TT, UK; Department of Clinical Laboratory Sciences, College of Applied Medical Sciences, King Khalid University, Abha 62521, Saudi Arabia; Department of Internal Medicine, Carver College of Medicine, University of Iowa, Iowa City, IA 52242, USA; Division of Hematology and Hemostaseology, Department of Medicine I, Medical University of Vienna, Vienna, Austria; Department of Inflammation and Ageing, University of Birmingham, Birmingham, UK; ImmunoPharmaLab, Department of Pharmacy, School of Medicine and Surgery, University of Naples Federico II, Via Domenico Montesano 49, Naples 80131, Italy

**Keywords:** galectin-9, venous thrombosis, platelets, neutrophil extracellular traps, thrombo-inflammation, Thrombosis, Vascular Biology, Inflammation, Basic Science Research

## Abstract

**Background:** Venous thromboembolism is increasingly recognized as a thrombo-inflammatory disease, yet the pathways linking inflammation to venous thrombus initiation remain incompletely defined. Galectin-9 (Gal-9), a β-galactoside-binding lectin elevated in inflammatory disorders, has no defined role in venous thrombosis.

**Methods:** Using plasma from patients with deep vein thrombosis (DVT), Gal-9-deficient mice, an inferior vena cava stenosis model, intravital microscopy, microfluidics, and ex vivo platelet and neutrophil assays, we assessed the functional contribution of Gal-9 to venous thrombosis.

**Results:** Plasma Gal-9 was elevated in DVT patients, correlated with D-dimer at presentation, and declined over 3-month follow-up. Gal-9 deletion did not alter hemostasis but markedly impaired thrombus initiation, with reduced platelet and neutrophil recruitment to the vessel wall and reduced intra-thrombus neutrophil extracellular trap (NET) formation. TNF-α/IFN-γ-stimulated endothelial cells upregulated Gal-9 and promoted platelet adhesion and fibrin(ogen) deposition, which Gal-9 neutralization partially reversed. Under venous shear, immobilized recombinant Gal-9 potentiated platelet activation and thrombus formation, particularly alongside collagen, whereas Gal-9 inhibition suppressed these responses. Gal-9-activated platelets promoted NETosis and shifted thrombus composition toward a NET-rich inflammatory phenotype.

**Conclusions:** Gal-9 is a previously unrecognized driver of venous thrombo-inflammation, linking vascular inflammation to platelet activation, NET formation, and thrombosis. Genetic deletion attenuated these responses in vivo, and antibody-mediated Gal-9 neutralization suppressed platelet activation and thrombus formation in vitro, identifying the Gal-9 glycan-binding pathway as a therapeutic target that disrupts thrombosis without impairing hemostasis.

## Introduction

Venous thromboembolism (VTE), comprising deep vein thrombosis (DVT) and pulmonary embolism, is a leading cause of preventable death. Yet it is still largely treated as a disorder of coagulation rather than of inflammation. This view is changing: DVT is now understood as a thrombo-inflammatory process in which activated endothelium, platelets, and innate immune cells cooperate to initiate and propagate the thrombus.^1,2^

Unlike arterial thrombosis, DVT develops at a morphologically intact vessel wall in the absence of plaque rupture or endothelial denudation. Reduced flow at venous valve sinuses generates local hypoxia and endothelial activation, triggering the release of Weibel-Palade body contents, P-selectin and von Willebrand factor (vWF), which support the capture of platelets and myeloid cells at the venous wall.^3,4^ Recruited monocytes deliver tissue factor to initiate coagulation, while neutrophils undergo NETosis, and the resulting neutrophil extracellular traps (NETs) provide a procoagulant, platelet-adhesive scaffold that activates the intrinsic pathway and stabilizes the growing thrombus.^2,3,5^ DVT is therefore initiated and propagated through reciprocal interactions between activated endothelium, platelets and innate immune cells, but the endothelial and immune-derived signals that couple this inflammatory activation to platelet and neutrophil recruitment, however, remain incompletely defined.

Galectins are β-galactoside-binding lectins that regulate immune and vascular function by clustering glycosylated receptors and modulating cell signaling.^6^ Gal-9, a tandem-repeat galectin containing two distinct carbohydrate recognition domains connected by a short peptide linker,^7^ is widely expressed across immune cell populations^8^ and is strongly induced in endothelial cells by inflammatory stimuli, particularly IFN-γ and TNF-α.^9–11^ Circulating Gal-9 rises in inflammatory, autoimmune, and cardiovascular disease and tracks with disease activity, most robustly in severe COVID-19, where it correlates with D-dimer and CRP.^12,13^ Within the vasculature, Gal-9 mediates neutrophil capture and adhesion through CD44 and β_2-_integrins and promotes atherosclerotic plaque formation,^11,14^ placing it at the interface of inflammation and vascular injury. How this glycan-dependent signaling contributes to thrombosis in the venous system, however, is far less understood than the protein-protein interactions that classically drive platelet activation. We recently showed that Gal-9 is itself a platelet agonist, activating platelets through the (hem)ITAM-coupled receptors GPVI and CLEC-2.^15^ Whether Gal-9 contributes to venous thrombus formation *in vivo*, and whether it links endothelial inflammation to platelet-driven thrombosis, has not been tested.

We tested the hypothesis that Gal-9 amplifies venous thrombosis by coupling inflammatory endothelial activation to platelet and neutrophil responses. Using plasma from patients with acute DVT, *Lgals9*-deficient mice, a murine inferior vena cava stenosis model of DVT, whole-blood microfluidics, and mechanistic *ex vivo* assays, we examined the contribution of Gal-9 to venous thrombus formation and asked whether it represents a hemostasis-sparing therapeutic target.

## Methods

Detailed methods and the Major Resources Table are provided in the Supplemental Material.

### Ethical Approval

Ethical approval for whole blood collection from healthy volunteers was obtained from the University of Birmingham Internal Ethical Review (ERN_11-0175). Human umbilical cords were collected from Birmingham Women’s and Sandwell and West Birmingham Hospitals NHS Trusts under ethical approval (REC 15/NE/0285) with written informed consent. All animal studies complied with the UK Animals (Scientific Procedures) Act 1986 and were approved by the UK Home Office (PPL PP8612246). DVT patient blood samples and blood samples from age- and sex-matched healthy controls were collected under ethical approval from the Medical University of Vienna (EK 1045/2020 and EK 039/2006).

### Patients sample analysis

Citrate-anticoagulated venous blood was collected from patients with acute DVT at baseline, i.e., within 21 days of diagnosis, and at 3-month follow-up within the framework of the prospective, observational BACH-VTE study.^16^ Additionally, citrate-anticoagulated venous blood was collected from sex- and age-matched healthy controls. D-dimer concentrations in patients with DVT were measured in the central laboratory of the Medical University of Vienna by a quantitative assay using a latex agglutination method, with a LLOQ of 0.27 mg/L. Platelet-poor plasma was isolated by centrifugation (3,450 × *g*, 15 min, room temperature) and stored at −80°C in the biobank facilities of the Medical University of Vienna in an ISO 9001:2015-certified environment until analysis.^17^ Plasma Gal-9 concentrations were measured by Luminex (R&D Systems) per the manufacturer’s instructions. The Gal-9/D-dimer association was assessed as described in the supplemental data.

### Murine DVT model: Inferior vena cava (IVC) stenosis

Male and female mice aged 8–10 weeks, all on a C57BL/6 background, were used for all experiments. Wild-type (WT) C57BL/6 mice were purchased from Jackson Laboratories. Gal-9-deficient mice [B6(FVB)-*Lgals9^tm^*^1^*^.1Cfg^*/Mmucd; RRID:MMRRC_031952-UCD] were obtained from the Mutant Mouse Resource and Research Center (MMRRC) at the University of California, Davis, an NIH-funded strain repository.

DVT was induced by partial flow restriction of the inferior vena cava (IVC), adapted from a previously described protocol.^18^ Mice were anesthetized with isoflurane (5%) and maintained under continuous anesthesia in the supine position throughout. Following a midline laparotomy, the IVC was exposed, separated from the aorta, and side branches ligated with 7-0 polypropylene suture. Partial flow restriction was achieved by ligating the IVC around a 30-gauge needle spacer with 7-0 polypropylene suture, then removing the spacer to re-establish a fixed, reduced luminal diameter. The abdominal wall and skin were closed in layers with 6-0 suture. At 48 hours, blood was collected by retro-orbital bleeding under isoflurane anesthesia, mice were euthanized by cervical dislocation, and thrombi were harvested and weighed.

### Intravital microscopy

Platelet and neutrophil recruitment to the vessel wall was imaged in the stenosed IVC 3 hours after flow restriction. Arterial thrombus formation was recorded in real time following laser injury of cremaster arterioles, with circulating platelets labeled via jugular vein cannulation. Full details are in the supplemental methods.

### *In vitro* platelet, neutrophil, and thrombus assays

Washed human platelets were prepared from citrated whole blood, and platelet activation was assessed by flow cytometry. Whole-blood thrombus formation under flow was assessed by perfusion over microspots and endothelial cells in the Maastricht flow chamber, and whole-blood clot kinetics were measured by thromboelastography. NET formation within thrombi was quantified by immunofluorescence staining for myeloperoxidase (MPO) and citrullinated histone H3 (CitH3). NETosis in isolated human neutrophils was assessed by MPO, CitH3, and Hoechst immunofluorescence staining. Full details are provided in the supplemental methods.

### Data Analysis

All statistical analyses were performed using Prism 10 (GraphPad Software Inc.), and unless otherwise stated, data are expressed as mean ± SEM. Plasma Gal-9 and D-dimer concentrations (Figure 1) were analyzed using linear mixed-effects models; full details are provided in the supplemental methods. Statistical differences between two groups were assessed using an unpaired t-test or, for non-normally distributed data, the Mann-Whitney U test. Comparisons across multiple groups were performed using one-way or two-way analysis of variance (ANOVA) with multiple comparisons, or the Kruskal-Wallis test with multiple comparisons for non-normally distributed data, as specified in the figure legends. A *P* value of less than 0.05 was considered statistically significant.

**Figure 1.**
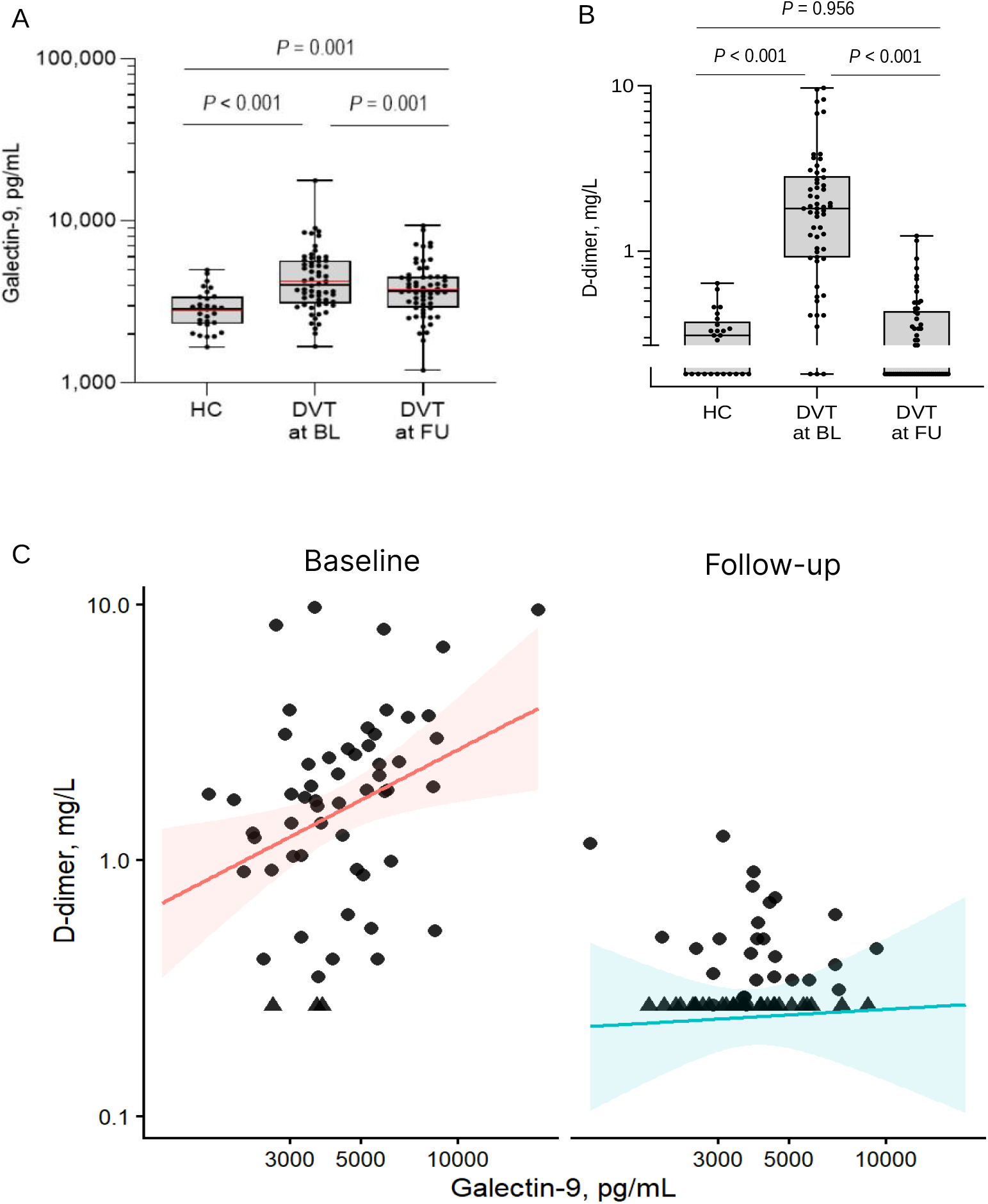
Plasma Gal-9 is elevated in DVT patients and decreased in follow-up. (A) Plasma galectin-9 concentration in healthy controls (HC; n = 30) and in patients with deep vein thrombosis (DVT) at baseline (BL; n = 60) and at 3-month follow-up (FU; n = 59). Data are shown on a logarithmic scale as box plots. Boxes span the lower and upper quartiles, whiskers span the minimum and the maximum values, horizontal lines within the boxes denote the median, and dots represent individual values. Data were analyzed using a linear mixed-effects model fitted to the log-transformed concentrations, with group–time category as a fixed effect and subject identifier as a random intercept. The displayed *P*-values are from model-based pairwise comparisons with Holm adjustment. (B) Plasma D-dimer concentration in HC (n = 25) and patients with DVT at BL (n = 58) and FU (n = 58). Data are shown on a logarithmic scale as box plots, as defined in panel A. The values below the lower limit of quantification (LLOQ; 0.27 mg/L) are shown in the lower segment of the *y*-axis. Data were analyzed using a left-censored linear mixed-effects model fitted to the log-transformed concentrations, with group–time category as a fixed effect and subject identifier as a random intercept. The displayed *P*-values are from model-based pairwise comparisons with Holm adjustment. (C) Association between galectin-9 and D-dimer concentrations in patients with DVT at BL and FU (n = 57 patients, each with both galectin-9 and D-dimer values available at BL and FU). Data are shown on logarithmic scales. Dots indicate individual D-dimer values. The values below the LLOQ are shown at the LLOQ. Data were analyzed using a left-censored linear mixed-effects model with log-transformed D-dimer concentration as the dependent variable and log-transformed galectin-9 concentration, time point, and their interaction as fixed effects, and subject identifier as a random intercept. Lines represent model-estimated geometric mean D-dimer concentrations across the observed range of galectin-9 concentrations, and shaded areas represent 95% confidence intervals for the estimated geometric means.

## Results

### Soluble Gal-9 is elevated in acute DVT

To assess the clinical relevance of Gal-9 in venous thrombosis, plasma Gal-9 levels were measured in patients with acute DVT at baseline and at 3-month follow-up and compared with healthy controls. Plasma Gal-9 was significantly elevated at baseline (n = 60) compared with controls (**Figure 1A, Table S1**). At follow-up (n = 59), Gal-9 levels were significantly reduced relative to baseline but remained higher than in controls (**Figure 1A**). At baseline, DVT patients also displayed significantly higher D-dimer, which normalized in most patients at follow-up (**Figure 1B**). A left-censored linear mixed-effects model revealed a significant positive correlation between Gal-9 levels with D-dimer levels at baseline but not at follow-up (**Figure 1C**).

These findings indicate that soluble Gal-9 levels are elevated during acute thrombosis and decline in the following months although remaining higher than healthy control, and can represent an additional thrombo-inflammatory biomarker for the acute phase of DVT.

### *In vivo* Gal-9 deficiency mitigates thrombus burden in a murine model of IVC stenosis

To establish the *in vivo* relevance of our findings, we assessed thrombus formation in Gal-9-deficient mice. *Lgals9*^-/-^mice were subjected to inferior vena cava (IVC) stenosis-induced venous thrombosis. Prior to thrombosis studies, hemostatic function was evaluated using a tail-bleeding assay, which revealed no significant difference in blood loss between WT and *Lgals9*^-/-^mice (**Figure 2A**), indicating preserved hemostatic function. Baseline platelet counts were comparable between genotypes, whereas neutrophil counts were modestly increased *in Lgals9*^-/-^mice (**Figure 2B-C**).

**Figure 2.**
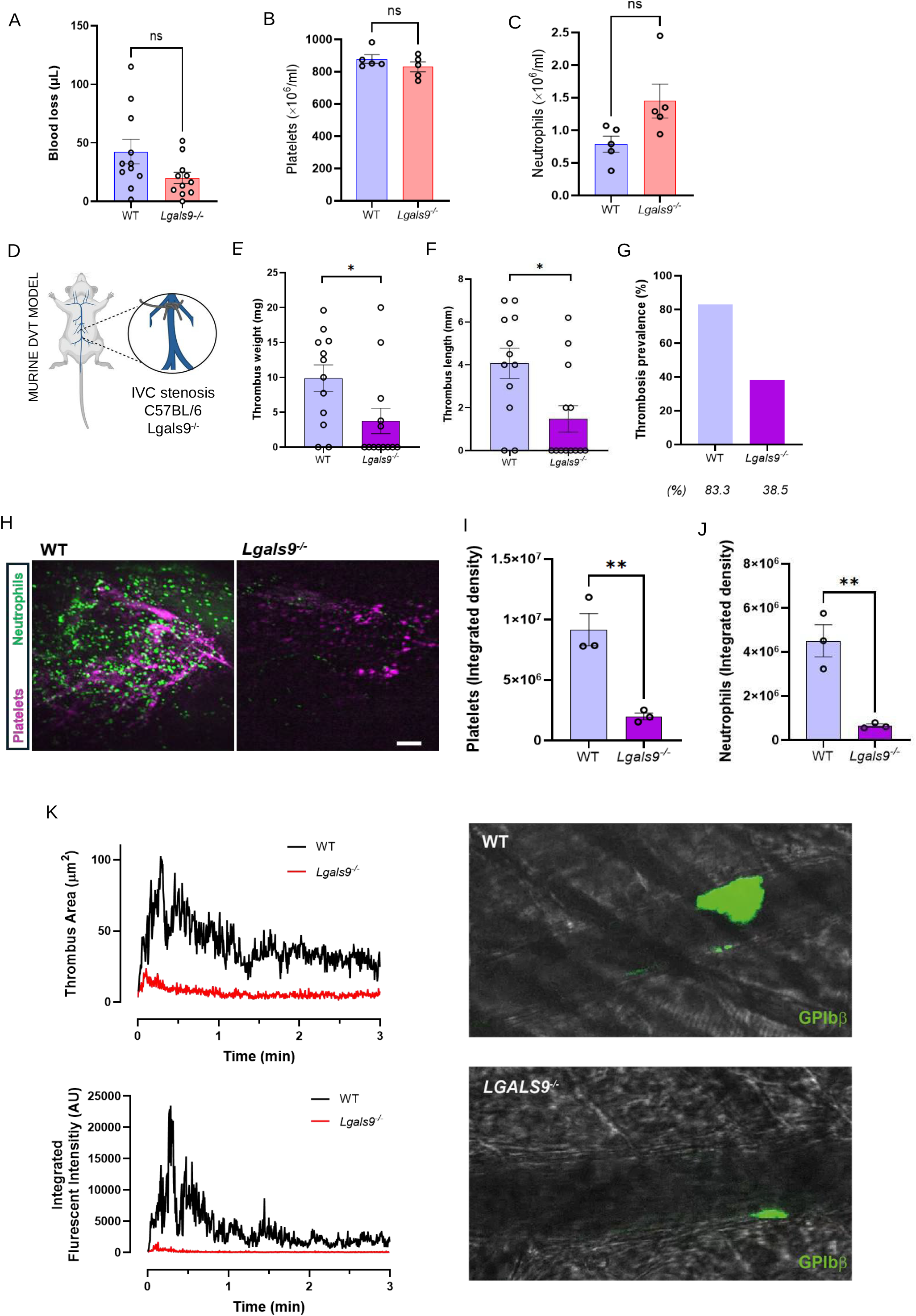
Genetic deletion of Gal-9 mitigates venous thrombosis in mice. (A) Blood loss (µL) measured by tail-bleed assay in WT (WT, n = 11) and Gal-9-deficient (*Lgals9*^-/-^, n = 9) mice. Mann-Whitney *U*-test, 2-tailed. (B) Circulating platelet count (×10⁶/mL) and (C) neutrophil count (×10⁶/mL) in WT and *Lgals9*^-/-^mice at baseline (n = 5 per group). Student *t*-test, 2-tailed, unpaired. (D) Schematic of the murine deep vein thrombosis (DVT) model: IVC stenosis was induced in WT and *Lgals9*^-/-^mice. (E) Thrombus weight (mg) and (F) thrombus length (mm) 48 hours after IVC stenosis (WT, n = 12 [3 female, 9 males]; *Lgals9*^-/-^, n = 13 [3 female, 10 male]). (G) Thrombosis prevalence (%) in WT (83.3%) versus *Lgals9*^-/-^(38.5%) mice. Mann-Whitney U-test 2-tailed. (H) Representative intravital images of the stenosed IVC in WT and *Lgals9*^-/-^mice showing platelets (magenta) and neutrophils (green). Scale bar = 100 µm. (I) Platelet and (J) neutrophil recruitment, quantified as integrated fluorescence density, in WT *versus Lgals9*^-/-^mice (n = 3 each group). Student *t*-test, 2-tailed, unpaired. (K) Laser-induced cremaster artery thrombus labeled by fluorescent *GPIbβ* monoclonal antibody in mice (n = 4 each group) with thrombus area and fluorescent intensity quantified. Results are expressed as mean ± SEM. Statistical analyses were done using either non-parametric Mann-Whitney U test or unpaired *t-test*. \**P* < .05, \*\**P* < .01, \*\*\*\**P* < .0001; ns, not significant.

Following 48 hours of IVC stenosis, *Lgals9*^-/-^mice exhibited a marked reduction in thrombus burden compared with WT controls. Thrombi were significantly smaller, with reduced weight and length, and overall thrombosis prevalence was significantly lower in *Lgals9*^-/-^mice (**Figure 2D-G**).

Since *Lgals9*^-/-^mice developed significantly smaller thrombi, we next investigated whether Gal-9 promotes early thrombo-inflammatory events during venous thrombosis. Intravital microscopy performed 3 hours after IVC stenosis revealed a marked reduction in both platelet and neutrophil recruitment to the vessel wall in *Lgals9*^-/-^mice compared to WT controls, as quantified by integrated fluorescence density (**Figure 2H-J, Video S1**). These findings identify Gal-9 as an important regulator of early platelet and leukocyte accumulation during thrombus formation.

To complement the IVC stenosis model and enable real-time visualization of platelet thrombus formation, we performed intravital laser injury of cremaster arteries to assess the contribution of Gal-9 to platelet thrombus growth *in vivo*. This complementary approach provided dynamic, real-time validation of the platelet phenotype identified *in vitro* and in the IVC stenosis model, rather than modeling arterial thrombosis as a distinct disease process. Following laser-induced vascular injury, platelet accumulation at the site of injury was reduced in *Lgals9*^-/-^mice compared with wild-type controls. Gal-9 deficiency resulted in a lower platelet fluorescence signal and impaired thrombus growth over the course of the recording (**Figure 2K, Video S2**). These findings provide independent, real-time *in vivo* evidence that Gal-9 contributes to platelet recruitment and thrombus growth. Although the cremaster arterial laser-injury thrombosis model represents a different vascular bed and mechanism of injury from venous thrombosis, the results support the platelet-dependent mechanism underlying the reduced thrombotic phenotype observed in the IVC stenosis model.

These findings demonstrate that Gal-9 promotes thrombosis in both venous and arterial settings *in vivo*, and that its genetic deletion protects against thrombus formation without impairing hemostasis.

### Gal-9 deficiency reduces intra-thrombus NETosis *in vivo*

To determine whether Gal-9 localizes within the developing thrombus, co-immunofluorescence staining for CD41 and Gal-9 was performed. Gal-9 was abundantly detected throughout WT thrombi and was particularly enriched within platelet-rich regions (**Figure 3A**). Quantitative analysis confirmed significant reductions in both absolute CD41+ area (mm²) and CD41+ area relative to total thrombus size in *Lgals9*^-/-^mice (**Figure 3B-C**). In addition, thrombus sections were stained positively for fibrinogen and Gal-9 together (**Figure S3A**). Given the marked reduction in neutrophil recruitment, we next examined whether Gal-9 influences NET formation. Co-staining for MPO and CitH3 demonstrated extensive NETosis within WT thrombi, whereas *Lgals9*^-/-^thrombi exhibited substantially reduced MPO/CitH3 co-localization (**Figure 3D**). Quantitative analysis confirmed significant reductions in both absolute NET area (mm²) and NET area relative to total thrombus size in *Lgals9*^-/-^mice (**Figure 3E-F**).

**Figure 3.**
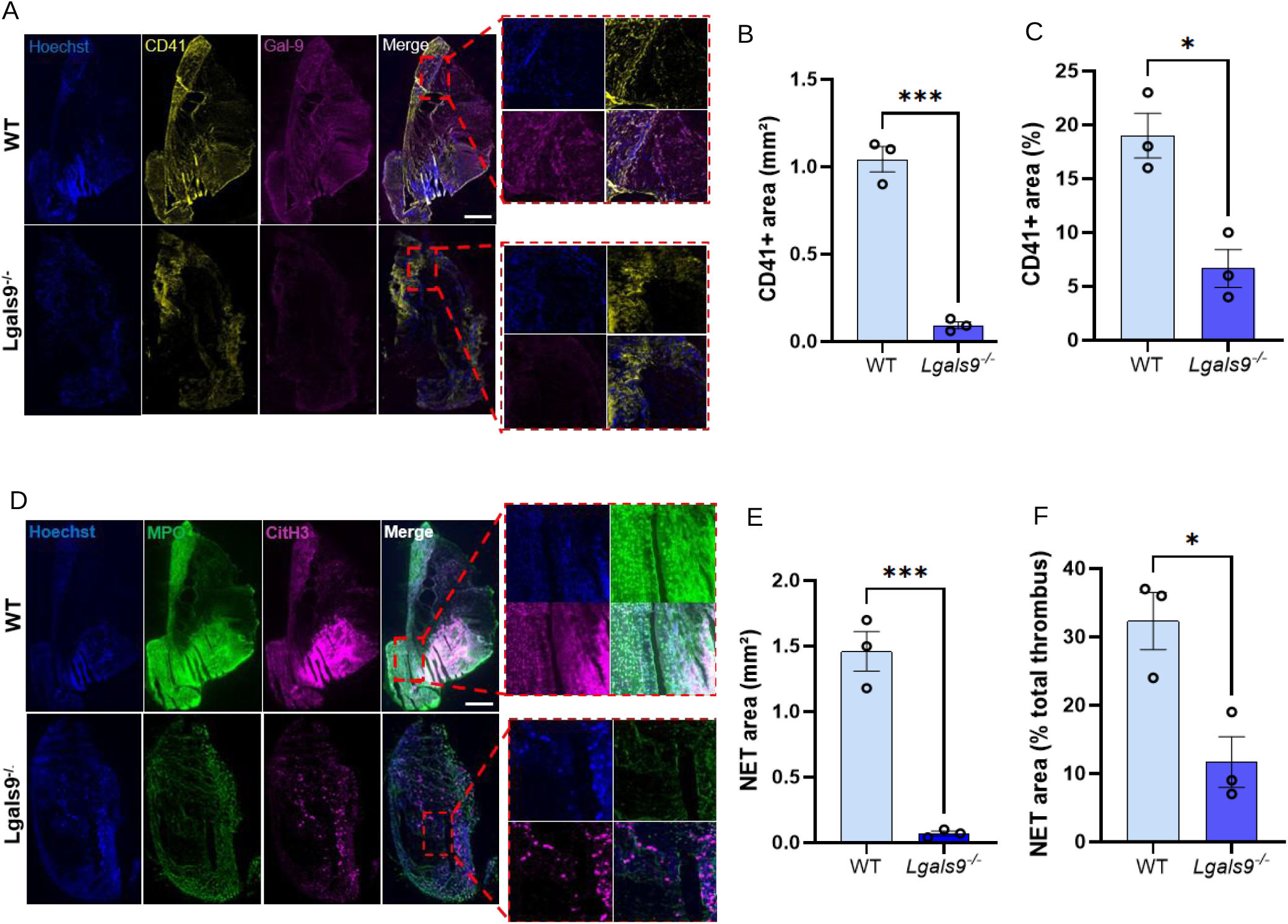
Genetic deletion of Gal-9 reduces NETosis within the venous thrombus *in vivo*. (A) Representative immunofluorescence images of thrombus cryosections co-stained for Hoechst (blue), CD41 (platelets, yellow), and Gal-9 (magenta), with merged channels, in WT (top) and *Lgals9*^-/-^(bottom) mice, showing Gal-9 localization within the platelet-rich thrombus. Scale bar = 500 µm. (B) CD41+ platelet area was quantified as absolute area (mm²) of total thrombus cross-sectional area, as well as (C) relative percentage of total thrombus area, in WT versus *Lgals9*^-/-^thrombi (n = 3 per group). Student *t*-test, 2-tailed, unpaired. (D) Representative immunofluorescence images of thrombus cryosections co-stained for Hoechst (blue), MPO (green), and CitH3 (magenta), with merged channels, in WT (top) versus *Lgals9*^-/-^(bottom) mice. Scale bar = 500 µm. (E) NET area (mm²) and (F) NET area as a percentage of total thrombus cross-sectional area, quantified from MPO/CitH3 co-positive staining in WT versus *Lgals9*^-/-^thrombi (n = 3 per group). Student *t*-test, 2-tailed, unpaired. Data are mean ± SEM. \**P* < .05, \*\**P* < .01, \*\*\*\**P* < .0001.

Collectively, these findings establish Gal-9 as a central amplifier of thrombo-inflammation by promoting platelet, neutrophil recruitment and NET formation within the developing thrombus. These data provide mechanistic evidence that Gal-9 orchestrates cellular interactions that sustain thrombus propagation during venous thrombosis.

### Endothelial activation-driven venous thrombosis is partially mediated by direct interaction of blood cells with Gal-9

As endothelial cells represent one of the major sources of Gal-9 during inflammation,^11,14,19^ we assessed whether inflammatory activation enhances endogenous Gal-9 expression. Endothelial cells stimulated with TNF-α and IFN-γ exhibited significantly increased Gal-9 expression compared with TNF-α alone, as demonstrated by flow cytometry (**Figure 4A**) and immunofluorescence (**Figure 4C**). ICAM-1 expression was robustly upregulated by TNF-α and further enhanced by combined TNF-α and IFN-γ stimulation (**Figure 4B**), confirming effective endothelial activation.

**Figure 4.**
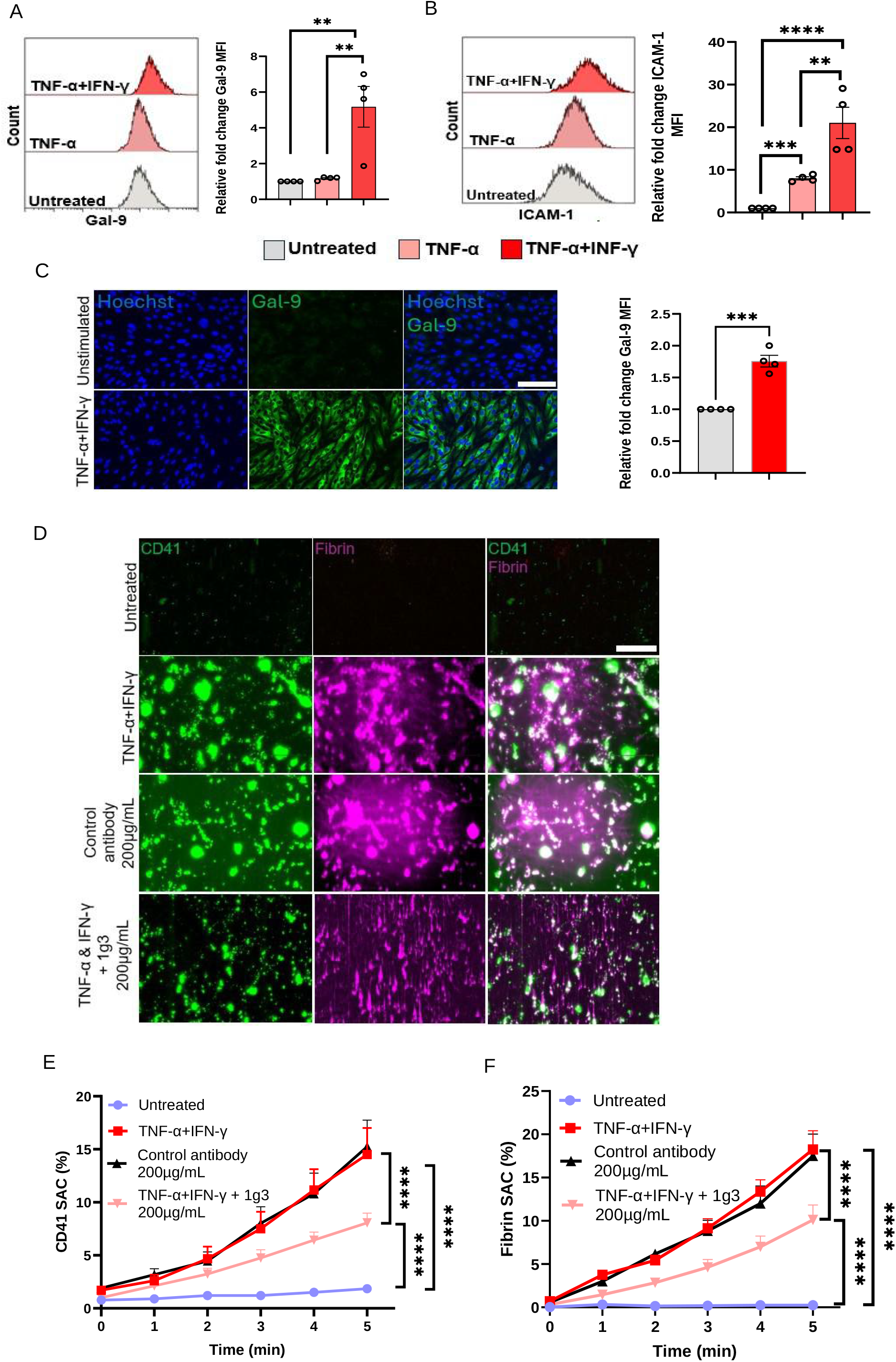
Endothelial Gal-9 directly potentiates platelet thrombus formation. (A) Gal-9 and (B) ICAM-1 surface expression on human umbilical vein endothelial cells (HUVECs) measured by flow cytometry following stimulation with TNF-α (100 U/mL) alone or TNF-α combined with IFN-γ (20 ng/mL) for 24 hours; representative histograms (left) and relative fold change in mean fluorescence intensity (MFI, right) versus untreated controls (n = 4). Ordinary one-way ANOVA. (C) Representative immunofluorescence images (Hoechst, blue; Gal-9, green) and quantification of relative Gal-9 MFI in unstimulated versus TNF-α/IFN-γ-stimulated HUVECs (n = 4). Scale bar = 100 µm. Student *t*-test, 2-tailed, unpaired. (D) Recalcified whole blood was perfused at venous shear rate (100 s⁻¹) over untreated or TNF-α/IFN-γ-stimulated HUVECs, in the presence or absence of the Gal-9-neutralizing antibody (clone 1g3, 200 µg/mL); representative images of platelet adhesion (CD41, green) and fibrin(ogen) deposition (magenta) at 5 minutes, with merged channels. Scale bar = 100 µm. (E) Platelet surface area coverage (SAC, %) and (F) fibrin SAC (%) quantified in real time over 5 minutes for untreated, TNF-α/IFN-γ-stimulated, and TNF-α/IFN-γ-stimulated + 1g3 conditions (n = 4). Two-way ANOVA with Tukey’s multiple comparison test. Data are mean ± SEM. **P < .01, ***P < .001, ****P < .0001.

To determine the functional consequence of increased endothelial Gal-9 expression, partially recalcified whole blood was perfused over cytokine-stimulated endothelial cells under venous shear. This resulted in significantly increased platelet adhesion and fibrin(ogen) deposition compared with untreated endothelium (**Figure 4D-F**). To confirm that this prothrombotic effect is specifically attributable to Gal-9, endothelial cells were treated with an inhibitory anti-Gal-9 monoclonal antibody (1g3 clone), which markedly attenuated both platelet adhesion and fibrinogen deposition over stimulated endothelial cells (**Figure 4D-F**). TNF-α and IFN-γ therefore upregulate endothelial Gal-9, creating a prothrombotic surface that supports platelet recruitment and fibrinogen deposition.

### Gal-9 potentiates platelet activation and thrombus formation under flow

Given the elevated Gal-9 levels observed in DVT and our prior identification of Gal-9 as a platelet agonist acting via GPVI and CLEC-2,^15^ we assessed whether Gal-9 directly promoted procoagulant platelet activation. In accordance with our previous data, stimulation of washed human platelets with recombinant Gal-9 induced dose dependent platelet activation, as evidenced by increased integrin GPIIb/IIIa activation, P-selectin expression, and phosphatidylserine externalization (**Figure S1A-C**).

In addition, Gal-9 significantly potentiated thrombin-induced platelet aggregation (**Figure S1D**), indicating that Gal-9 amplifies responses to classical platelet agonists. Furthermore, Gal-9 stimulation increased Ca^2+^ mobilization in a concentration-dependent manner (**Figure S1E**), consistent with engagement of GPVI and CLEC-2 dependent signaling pathways and a procoagulant platelet phenotype.

Together, these findings extend our previous observations and demonstrate that Gal-9 not only directly activates platelets but also enhances agonist-driven platelet responses.

Thrombus formation on immobilized Gal-9 under low shear conditions induced minimal platelet adhesion and activation (**Figure 5A-D**). In contrast, co-immobilization of Gal-9 with collagen markedly enhanced thrombus formation over time, as demonstrated by increased fibrinogen binding, P-selectin expression, and phosphatidylserine externalization (annexin V binding) compared with collagen alone (**Figure 5A-D**). Combining Gal-9 with von Willebrand factor (vWF) did not produce a comparable increase in thrombus formation, although a significant increase in P-selectin expression was observed (**Figure 5A-D**). In contrast, at arterial shear rate, Gal-9 co-immobilized with collagen or vWF did not produce a comparable enhancement of platelet activation or thrombus formation *in vitro* (**Figure S2**).

**Figure 5.**
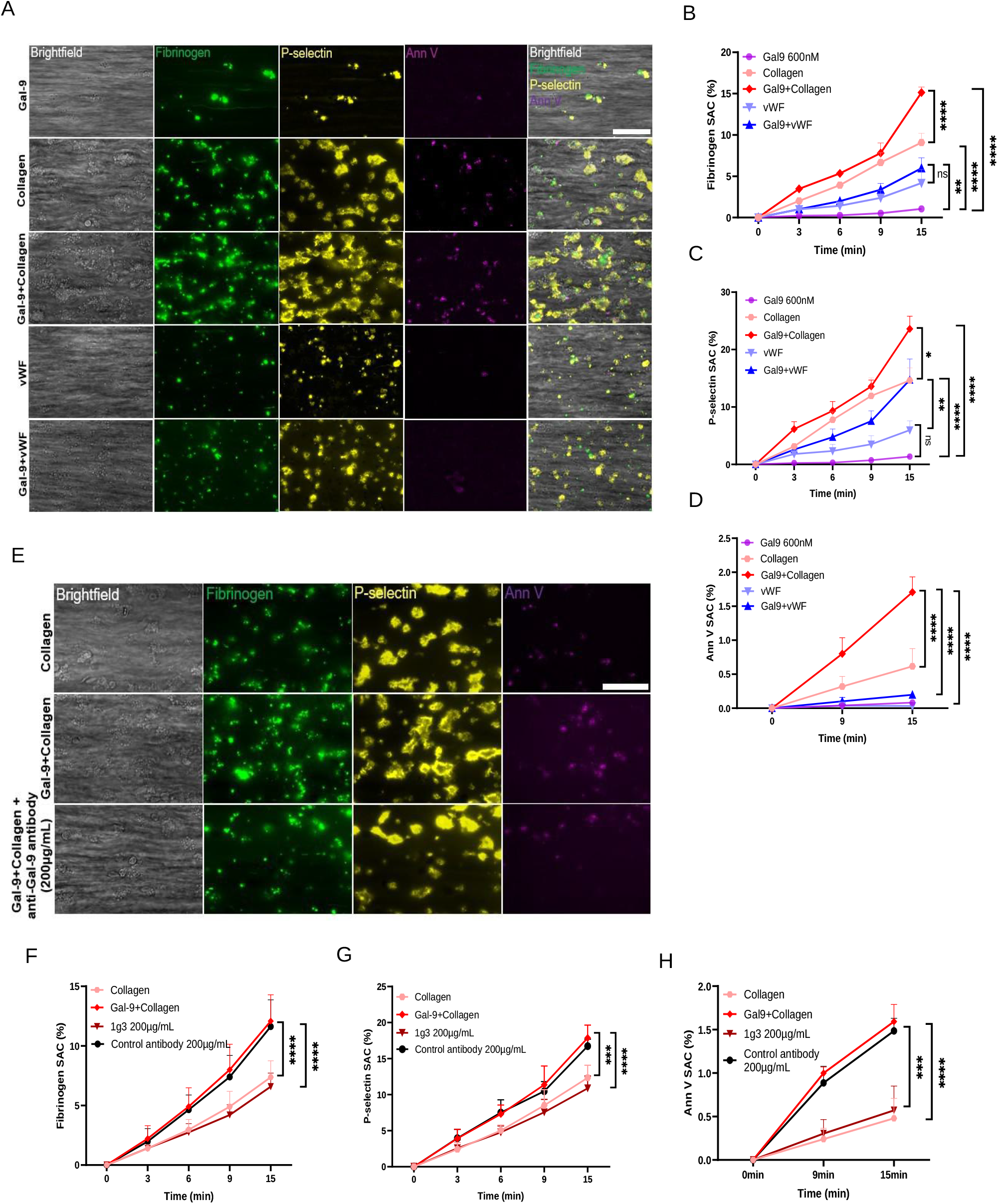
Gal-9 accelerates platelet thrombus formation at venous shear rate *in vitro*. (A) Recalcified whole blood was perfused at venous shear rate (100 s⁻¹) over microspots coated with immobilized Gal-9 (600 nM), collagen (50 µg/mL), von Willebrand factor (vWF, 100 µg/mL), or combinations thereof (Gal-9 + collagen; Gal-9 + vWF). Platelet activation was assessed by fibrinogen binding (AF488), P-selectin expression (PE), and phosphatidylserine exposure (annexin V-AF647); representative multichannel images at 15 minutes. Scale bar = 50 µm. (B) Fibrinogen SAC (%), (C) P-selectin SAC (%), and (D) annexin V SAC (%) quantified over 15 minutes (n = 4). Two-way ANOVA with Tukey’s multiple comparison test. (E) Representative images of whole blood perfused over collagen alone, Gal-9 + collagen, or Gal-9 + collagen in the presence of the Gal-9-neutralizing antibody (clone 1g3, 200 µg/mL). Scale bar = 50 µm. (F) Fibrinogen SAC (%), (G) P-selectin SAC (%), and (H) annexin V SAC (%) quantified over 15 minutes for collagen, Gal-9 + collagen, Gal-9 + collagen + 1g3, and Gal-9 + collagen + isotype control antibody (200 µg/mL) (n = 3). Two-way ANOVA with Tukey’s multiple comparison test. Data are mean ± SEM. \**P* < .05, \*\**P* < .01, \*\*\*\**P* < .0001; ns, not significant.

To determine whether this effect at venous shear was specifically mediated by Gal-9, whole blood was perfused over Gal-9 and collagen-coated surfaces in the presence of the Gal-9 blocking 1g3 antibody. Inhibition of Gal-9 significantly reduced platelet activation and thrombus formation, as reflected by decreased fibrinogen binding, P-selectin expression, and decreased phosphatidylserine exposure (**Figure 5E-H**). Together, these findings demonstrate that Gal-9 acts in concert with collagen to potentiate platelet activation and thrombus formation under venous shear conditions.

To model the impact of circulating Gal-9, we tested whether soluble Gal-9 enhances platelet-driven thrombus formation under flow. Whole blood supplemented with recombinant Gal-9 was perfused over collagen-coated surfaces at venous shear.

Soluble Gal-9 enhanced platelet-mediated thrombus formation in a concentration-dependent manner (**Figure 6A**), as shown by increased fibrinogen binding, P-selectin expression, and phosphatidylserine externalization (**Figure 6B-G**). These findings indicate that soluble Gal-9, similar to immobilized Gal-9, potentiates platelet activation under flow conditions.

**Figure 6.**
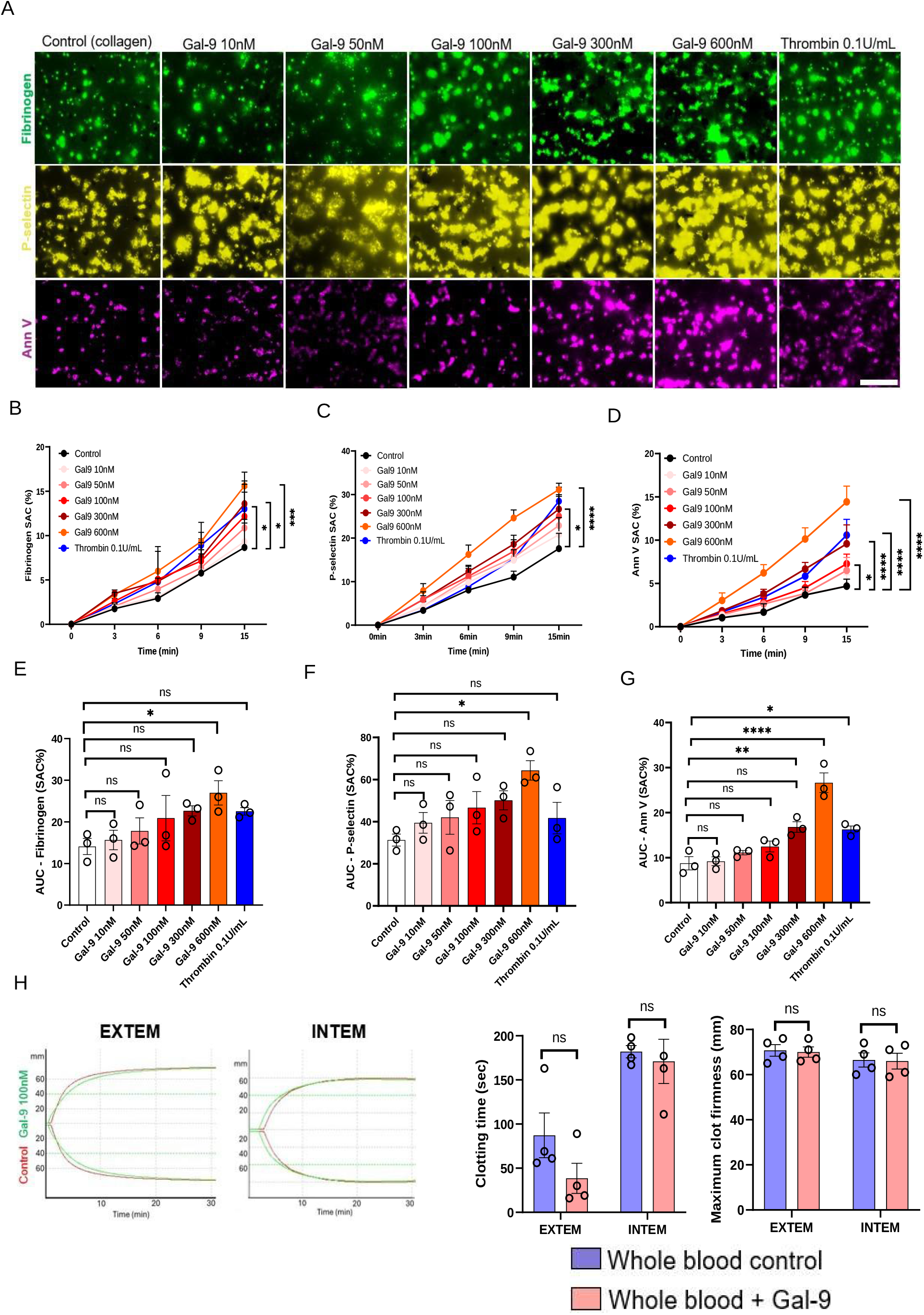
Gal-9 directly potentiates whole blood thrombus formation. (A) Recalcified whole blood spiked with increasing concentrations of recombinant Gal-9 (10, 50, 100, 300, 600 nM), thrombin (0.1 U/mL, positive control), or vehicle (collagen alone, control) was perfused over immobilized collagen (50 µg/mL) at venous shear rate (100 s⁻¹). Representative images of fibrinogen binding (AF488), P-selectin expression (PE), and phosphatidylserine exposure (annexin V-AF647) at 15 minutes. Scale bar, 50 µm. (B) Fibrinogen SAC (%), (C) P-selectin SAC (%), and (D) annexin V SAC (%) quantified over 15 minutes across all conditions (n = 3). Two-way ANOVA with Tukey’s multiple comparison test. (E–G) Area under the curve (AUC) for fibrinogen SAC, P-selectin SAC, and annexin V SAC, respectively, across Gal-9 concentrations and thrombin control. One-way ANOVA with Tukey’s multiple comparison test. (H) Thromboelastography (ROTEM) of whole blood ± Gal-9 (100 nM): representative EXTEM and INTEM traces (left), clotting time (seconds) and maximum clot firmness (mm) (right) (n = 4). Student *t*-test, 2-tailed, unpaired. Data are mean ± SEM. *P < .05, **P < .01, ****P < .0001; ns, not significant.

To determine whether Gal-9 also modulates coagulation at the whole blood level, thromboelastography was performed. Addition of Gal-9 reduced clotting time without affecting clot firmness (**Figure 6H**), indicating accelerated coagulation kinetics without altering clot formation. Together, these findings demonstrate that both immobilized and soluble Gal-9 enhances platelet-driven thrombus formation and promotes a procoagulant state and supports thrombosis suggesting that soluble Gal-9 is not merely a biomarker of cell activation but also a direct contributor of thrombo-inflammation.

### Gal-9 activated platelets amplify NET formation

Given the established role of NETs in venous thrombosis,^20,21^ and our previous findings that Gal-9 directly activates neutrophils,^11^ we next assessed whether Gal-9 promotes NETosis directly and/or platelet-dependent NET formation. Recombinant Gal-9, unstimulated platelets, or Gal-9-stimulated platelets were added to adherent neutrophils and NETosis was assessed by MPO, CitH3, and Hoechst immunofluorescence staining (**Figure 7A**). Gal-9 alone significantly increased NET formation compared with unstimulated neutrophils, as evidenced by increased MPO, CitH3, and extracellular Hoechst staining, with characteristic web-like structures observed by immunofluorescence (**Figure 7A & B**). Addition of unstimulated platelets did not induce NET formation, whereas Gal-9-activated platelets resulted in significantly greater NETosis than Gal-9 alone, with further increases in MPO, CitH3, and Hoechst staining (**Figure 7A & B**).

**Figure 7.**
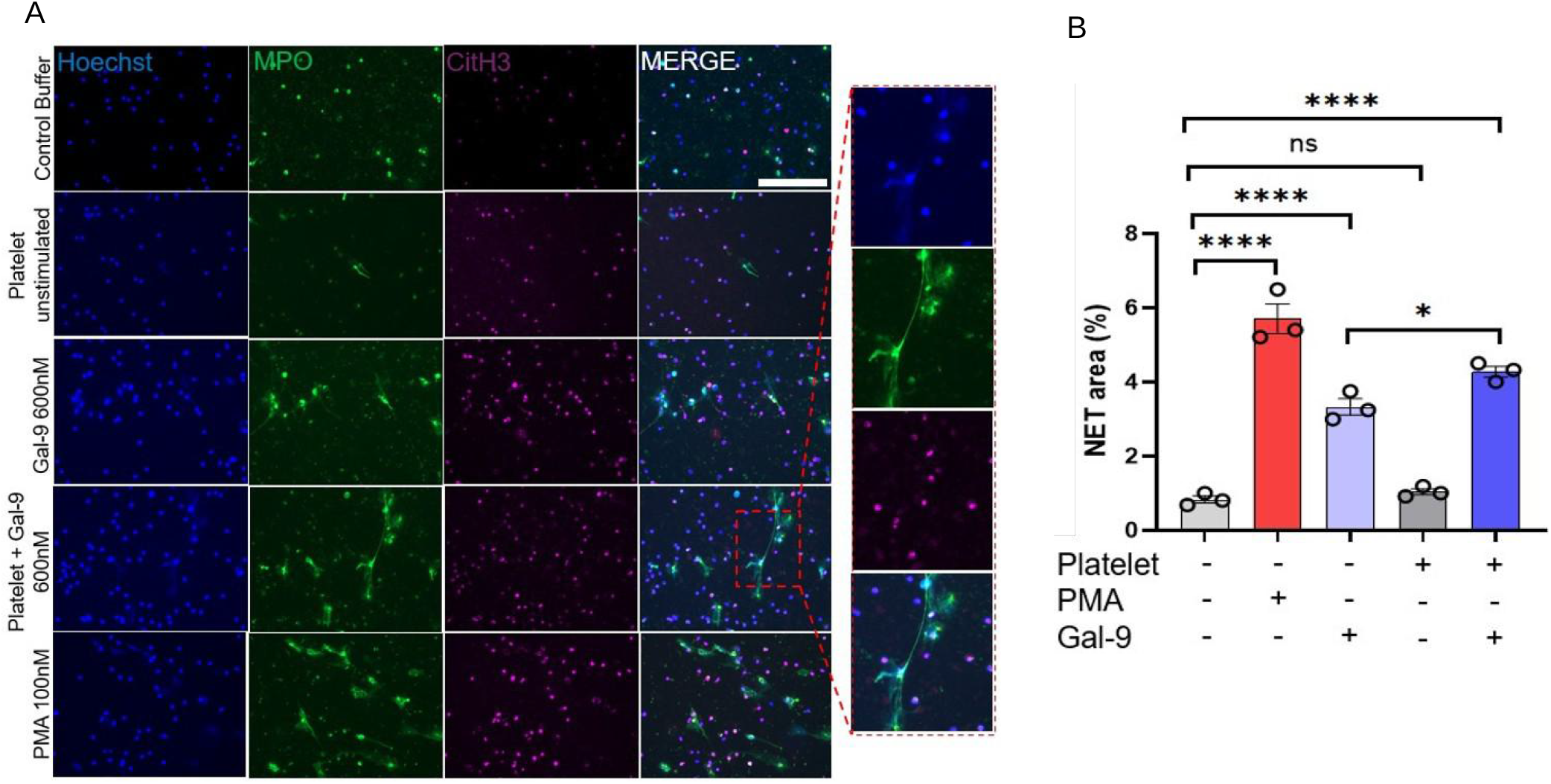
Gal-9 activated platelet induced neutrophil NETosis. (A) Representative immunofluorescence images (Hoechst, blue; MPO, green; CitH3, magenta; merge) of isolated neutrophils incubated for 3 hours with: buffer control; PMA (100 nM, positive control); Gal-9 alone (600 nM); platelets pre-activated with Gal-9 (600 nM); or platelets alone (unstimulated). Scale bar = 200 µm. (B) Quantification of neutrophil extracellular trap (NET) area as a percentage of total field area, averaged across 5 fields of view per condition (n = 3). Statistical significance was determined by one-way ANOVA with Tukey’s multiple comparisons test. Data are mean ± SEM. *P < .05, **P < .01, ****P < .0001; ns, not significant.

Together, these findings demonstrate that Gal-9 directly induces NET formation and that Gal-9 activated platelets further amplify this response, highlighting cooperation between platelets and neutrophils as an additional mechanism through which Gal-9 promotes thrombo-inflammation.

## Discussion

In this study, we show that Gal-9 is required for venous thrombus formation in mice, acts at the initiation of thrombosis rather than only at propagation, and does so without a measurable alteration in hemostasis. *Lgals9*-deficient mice formed fewer and smaller thrombi after IVC stenosis, recruited fewer platelets and neutrophils to the vein wall within 3 hours, and produced thrombi with less platelet and NET content. In parallel, TNF-α and IFN-γ-activated endothelium acquired Gal-9 and captured platelets in a partly Gal-9-dependent manner, and Gal-9 amplified collagen-driven platelet activation under venous shear. Finally, Gal-9-activated platelets drove NET formation more strongly than Gal-9 alone. These observations converge on Gal-9 as a mediator that couples inflammatory endothelial activation to platelet and neutrophil responses that drive the initiation of venous thrombosis.

Although anticoagulation remains the gold standard for VTE management, targeting inflammatory pathways to overcome anticoagulant resistance and reduce recurrence is increasingly recognized as a valuable adjunct. Even with direct fibrinolysis, anticoagulation often increases bleeding risk without reliably preventing recurrence, particularly in cancer-associated VTE,^22,23^ underscoring the need for strategies that better balance efficacy against bleeding risk. VTE pathogenesis reflects an interplay between coagulation and inflammation, immunothrombosis or thrombo-inflammation, such that inflammatory activity persisting after anticoagulation can drive recurrence. VEXAS syndrome, caused by somatic UBA1 mutation, illustrates this. VTE occurs in over 40% of patients, recurs in 41%, and develops despite therapeutic anticoagulation in 20%.^24^ Thrombotic events cluster with active inflammatory disease,^25,26^ and hyperactivated immune cells with heightened cytokine release drive endothelial dysfunction, further raising recurrence risk.^27^ Resolving thrombosis therefore likely requires mitigating inflammation alongside anticoagulation; targeting inflammatory mediators, whether alone or in combination with anticoagulants, may offer a more effective strategy for clearing treatment-resistant clot and preventing further thrombotic events.

The dissociation between Gal-9 and D-dimer kinetics over the disease course suggests that circulating Gal-9 does not simply track thrombus burden. Gal-9 remained elevated above control levels at follow-up despite normalization of D-dimer, and its baseline correlation with D-dimer was lost over time. Rather than resolving in parallel with the clot, Gal-9 remained raised, which is consistent with sustained vascular inflammation or endothelial activation after the acute event rather than with the acute thrombotic process itself. As our study lacks pre-thrombotic samples, we cannot determine whether the elevation precedes thrombosis and reflects a predisposing inflammatory state or arises as a consequence of it, and this distinction will require prospective sampling before an event. Whether Gal-9 adds information to D-dimer for diagnosis or risk stratification requires a prospective cohort with pre-event sampling and recurrence endpoints.^28–30^ Elevated circulating Gal-9 has been reported in other thrombo-inflammatory conditions, including severe COVID-19^12,13^ and large-artery atherosclerotic stroke,^31^ correlating with CRP and D-dimer, though these associations remain largely correlative, and the mouse data presented here are what argue for a causal contribution.

Our finding that Gal-9 deficiency reduces thrombus burden *in vivo* is consistent with its role in other inflammatory vascular diseases. We have previously shown that *ApoE*^-/-^*Lgals9*^-/-^mice develop significantly reduced aortic plaque burden compared with *ApoE*^-/-^controls, an effect attributed to Gal-9-driven monocyte recruitment and a pro-inflammatory macrophage phenotype.^14,32^ Similarly, TNF-α/IFN-γ-induced endothelial Gal-9 mediates neutrophil capture and adhesion via CD44 and β_2-_integrin, and neutrophil recruitment was significantly reduced in Gal-9-deficient mice in a model of zymosan-induced peritonitis, identifying endothelial-derived Gal-9 as a direct driver of leukocyte recruitment *in vivo*.^11^ A comparable pathogenic role has recently been described in neurodegenerative disease: microglia-derived Gal-9 generates neurotoxic amyloid-β fibrils, and Gal-9 knockout attenuates amyloid pathology, neuroinflammation, and cognitive impairment in APP/PS1 mice.^33^ Endothelial Gal-9 is also inducible and epigenetically regulated rather than constitutive, being controlled by histone deacetylase 3,^10^ which places it among the set of inducible endothelial effectors that reprogram the vein wall toward a proinflammatory, procoagulant state. However, the global knockout cannot attribute the phenotype to a cell type, and endothelial, platelet and myeloid conditional deletion will be needed to apportion it.

Gal-9 behaves in this setting like a secreted alarmin. Its actions on platelets via the ITAM receptors GPVI and CLEC-2,^15^ parallel those of HMGB1, which shortens clot formation time in thromboelastometry and drives NETosis through RAGE and TLR2,^34^ and S100A8/A9, which generates procoagulant platelets through GPIbα.^35^ Our data extend this parallel in one specific direction: the platelet is not just a target of Gal-9 but an amplifier of it, since Gal-9-activated platelets induced more NET formation than Gal-9 alone, and both platelet and NET content of the thrombus fell in *Lgals9*-deficient mice. That is mechanistically consistent with the report that Gal-9 upregulates PAD-4 and histone citrullination in granulocytes.^36^ Direct evidence that elevated Gal-9 is itself damaging, not merely a marker, comes from a study identifying a hyperactive Gal-9^high^ expressing neutrophil population that drives radiation-induced frailty:^37^ these cells showed elevated ROS, NETs, and IFN-γ, caused direct multi-organ injury, and resisted macrophage clearance via CD47. Critically, the relationship was causal and bidirectional, recombinant Gal-9 worsened frailty in healthy mice, and anti-Gal-9 antibody reversed it in irradiated mice, without detectable toxicity to endothelium, B, or T cells.^37^ The Gal-9 concentrations used *in vitro* exceed patient plasma levels, a limitation, although galectins accumulate locally through endothelial release, leukocyte secretion, vesicle trafficking and matrix retention, and Gal-3 reaches concentrations in the venous thrombus and vein wall far above circulating levels.^38^ Whether the same holds for Gal-9 requires direct measurement in thrombus tissue.

Persistent endothelial activation after DVT may underlie incomplete thrombus resolution and chronic venous complications despite adequate anticoagulation, as sustained adhesion-molecule elevation in patients with residual obstruction or post-thrombotic syndrome shows that vascular thrombo-inflammation outlasts the acute event.^39,40^ The endothelium is therefore an attractive target: an activated, proinflammatory, procoagulant endothelium drives platelet recruitment, leukocyte adhesion, and coagulation, and endothelial-directed therapy could offer greater spatial selectivity than systemic anticoagulants or antiplatelets. Blocking P-selectin, E-selectin, VCAM-1, or ICAM-1 has been proposed to limit leukocyte trafficking, but single-target blockade has repeatedly failed to achieve consistent efficacy, in part because these adhesion receptors compensate for one another.^41^ Endothelial cells are a well-established source of Gal-9 across vascular beds and disease models. IFN-γ stimulation induces Gal-9 expression in cultured human endothelial cells, which in turn enhances leukocyte adhesion to the endothelial monolayer, an effect abolished by lactose-mediated blockade of Gal-9 lectin activity.^9^ Although Gal-9 driven activation of inflammatory pathways has not been fully explored in thrombosis, studies in other settings show galectins actively amplify inflammatory signaling rather than simply marking it. Gal-9 enhances inflammatory cytokine gene expression in monocytes,^42^ potentiates TNF-α/IL-6 production by microglia.^43^ Related proinflammatory activity has been reported for Gal-3 (IL-6/CXCL8 induction)^44^ and Gal-8 (CXCL10 induction),^45^ and several galectins directly activate platelets and neutrophils through distinct receptors, Gal-1/-8 as direct platelet agonists,^46^ Gal-3 via Dectin-1,^47^ and Gal-8 via integrin α_M_β_2_-dependent neutrophil adhesion, with both Gal-8 and Gal-9 inducing oxidative burst.^48^ This is directly relevant to DVT pathogenesis, in which the endothelium is reprogrammed toward a proinflammatory, procoagulant phenotype, co-expressing tissue factor, adhesion molecules, and other mediators that together support platelet activation, leukocyte trafficking, and thrombus formation, placing Gal-9 among a coordinated set of inducible endothelial effectors rather than acting in isolation. Consistent with this, platelet adhesion and fibrin deposition on cytokine-activated, Gal-9-expressing endothelial cells were partially, but significantly, reduced by anti-Gal-9 inhibition in our data.

Beyond genetic knockouts, the prospect of galectin inhibition in the clinical setting appears promising: the Gal-3 inhibitor GB0139 was well tolerated in COVID-19 pneumonitis (DEFINE; NCT04473053),^49^ and belapectin has advanced for NASH cirrhosis (NCT02462967). For Gal-9, the humanized antibody LYT-200, in early-phase development (NCT04666688), shows preclinical efficacy in hematological malignancy,^50^ and has completed Phase 1b evaluation alone and with venetoclax/HMA in relapsed/refractory AML/MDS (NCT05829226),^51^ a population in which thrombosis is highly prevalent, with a reported prevalence of 22.1% in intensively treated AML/high-risk MDS.^52,53^ These trials included no thrombotic endpoints, but confirm Gal-9 blockade is clinically tractable and raise the possibility of antithrombotic benefit alongside antineoplastic effects, testable through exploratory endpoints nested in ongoing oncology trials.

In conclusion, we have demonstrated a major impact of Gal-9 in venous thrombosis. Gal-9 deficiency significantly reduced thrombus burden *in vivo*, with concomitant reductions in platelet and neutrophil recruitment and NETosis at the vein wall and targeting Gal-9 could potentially reduce thrombo-inflammatory complications.

## Acknowledgments

The authors gratefully acknowledge all patients and blood and umbilical cord donors for their participation in this study. The authors thank Dr Michael Feibig and Absolute Antibody for generously providing the Galectin-9 (Gal-9) neutralizing antibody (clone 1g3). The authors also thank the research nurses and midwifery teams at Birmingham Women’s and Children’s NHS Foundation Trust and Sandwell and West Birmingham Hospitals NHS Trust for their assistance with umbilical cord collection.

## Author Contributions

A.A.K.: conceptualization, methodology, investigation, data curation, formal analysis, writing – original draft, and writing – review & editing; A.A.M.: methodology and writing – review & editing; K.W.: methodology, visualization, and writing – review & editing; I.B.: formal analysis, visualization, and writing – review & editing; D.S.: methodology and writing – review & editing; C.A.: methodology and writing – review & editing; J.E.: methodology and writing – review & editing; D.L.: methodology, visualization, and writing – review & editing; H.M.: methodology, visualization, and writing – review & editing; D.K.: methodology, visualization, and writing – review & editing; F.M.: methodology, visualization, and resources; G.E.R.: conceptualization, funding acquisition, project administration, resources, supervision, and writing – review & editing; J.R.: conceptualization, funding acquisition, project administration, resources, supervision, and writing – review & editing; A.B.: conceptualization, funding acquisition, project administration, resources, supervision, and writing – review & editing; and A.J.I.: conceptualization, funding acquisition, project administration, resources, supervision, and writing – review & editing.

## Data Availability

The data that support the findings of this study are available from the corresponding author upon reasonable request. Detailed methods and supplemental data are provided in the Supplemental Material.

## Supplemental Material

Supplemental Methods

Figures S1–S3

Table S1 Videos S1–S2

Major Resources Table

References 54–56

## Novelty and Significance

### What Is Known?

- Venous thromboembolism is a thrombo-inflammatory disease in which activated endothelium, platelets, and neutrophils cooperate to initiate thrombosis, with neutrophil extracellular traps (NETs) forming a structural, procoagulant component of the thrombus.
- Galectin-9 is a β-galactoside-binding lectin that is elevated in inflammatory disease and regulates leukocyte trafficking and endothelial activation.
- Galectin-9 is itself a platelet agonist, activating platelets through the (hem)ITAM-coupled receptors GPVI and CLEC-2.

### What New Information Does This Article Contribute?

- Plasma galectin-9 is elevated in patients with acute deep vein thrombosis, correlates with D-dimer at presentation, and declines over 3-month follow-up.
- Galectin-9 deficiency impairs venous thrombus initiation, platelet and neutrophil recruitment to the vessel wall, and intra-thrombus NET formation without impairing hemostasis.
- Beyond direct agonism, galectin-9 potentiates platelet activation and thrombus formation under venous shear, enhances responses to classical agonists, and amplifies platelet-driven NETosis; antibody blockade reproduces the protection conferred by genetic deletion.

## Nonstandard Abbreviations and Acronyms

CitH3: citrullinated histone H3
CLEC-2: C-type lectin-like receptor 2
DVT: deep vein thrombosis
Gal-9: galectin-9
GPVI: glycoprotein VI
HUVEC: human umbilical vein endothelial cell
IVC: inferior vena cava
MPO: myeloperoxidase
NET: neutrophil extracellular trap
SAC: surface area coverage
VTE: venous thromboembolism
vWF: von Willebrand factor

