## Supplemental for "Galectin-9 drives thrombo-inflammatory circuits initiating venous thrombosis"

#### Table of Contents

Supplemental Methods

Table S1. Clinical characteristics of DVT patients

Major Resources Table

Figure S1. Gal-9 directly modulates platelet activation

Figure S2. Platelet thrombus formation at arterial shear rate in vitro

Figure S3. Co-immunostaining of fibrinogen and Gal-9 in murine thrombus

Video S1. IVC stenosis model of DVT

Video S2. Laser-injury arterial thrombosis

Supplemental References

### Supplemental Methods

#### Measurement of plasma galectin-9

Plasma galectin-9 concentrations were determined in platelet-poor plasma using a custom magnetic bead-based multiplex immunoassay (Human Luminex Discovery Assay, catalogue # LXSAHM; R&D Systems, Bio-Techne), in which galectin-9 was measured simultaneously with additional analytes in the same panel; only galectin-9 concentrations are reported here. The assay was performed according to the manufacturer’s instructions. Samples were thawed on ice immediately before analysis, and concentrations were interpolated from the analyte-specific standard curve supplied with the kit. [Co-measured analytes, sample dilution, and Luminex analyser model to be inserted.]

#### Statistical analysis of plasma Gal-9 and D-dimer concentrations

Plasma Gal-9 and D-dimer concentrations were analyzed using linear mixed-effects models fitted to log-transformed concentrations, with group–time category as a fixed effect and subject identifier as a random intercept, to account for repeated measures across baseline and follow-up. D-dimer values below the lower limit of quantification (LLOQ) were handled using a left-censored model. The association between Gal-9 and D-dimer concentrations was assessed using a left-censored linear mixed-effects model, with log-transformed D-dimer concentration as the dependent variable and log-transformed Gal-9 concentration, time point, and their interaction as fixed effects, and subject identifier as a random intercept. Pairwise comparisons were derived from the fitted models with Holm adjustment for multiple comparisons. All statistical analyses were performed in GraphPad Prism (version 10.2.3; GraphPad Software, Boston, MA, USA).

#### Intravital imaging of platelet and neutrophil recruitment in the stenosed IVC

Three hours after IVC stenosis, male mice (WT and *Lgals9^-/-^*, 8–10 weeks old) were anesthetized with intraperitoneal ketamine (76 mg/kg) and medetomidine (1 mg/kg) and injected retro-orbitally with DyLight 649-anti-GPIbβ/CD42c (X-649, Emfret Analytics; 2 µg) and PE-anti-Ly6G (BioLegend, clone 1A8; 4 µg) to label circulating platelets and neutrophils, respectively. The IVC was exposed and imaged supine on an Olympus BX61WI upright microscope with a 3i VIVO-SDC spinning-disk confocal unit (Yokogawa CSU-10, Photometrics Evolve EMCCD camera, 10× air objective), 1–2 mm caudal to the stenosing suture, for ≥1 minute per animal.

Videos were analyzed in Fiji/ImageJ. Platelet and neutrophil channels were split and analyzed separately. A fixed ROI over the vessel/thrombus, plus a background ROI, were applied uniformly across frames (with StackReg registration where drift was present), and integrated density and mean gray value were measured by Multi Measure. Background-corrected integrated density was calculated as vessel ROI integrated density minus (vessel ROI area × background mean intensity), giving final and maximum corrected values per animal for each channel.

#### Laser arterial injury

WT and *Lgals9^-/-^* mice (8–10 weeks old, 20–25 g, all male) were anesthetized intraperitoneally with a mix of ketamine (76 mg/kg) and medetomidine (1 mg/kg). Following tracheotomy, DyLight 649-conjugated anti-GPIbβ antibody (Emfret Analytics; 2 µg) was administered via jugular vein cannulation to label circulating platelets. The cremaster muscle was surgically exposed, and thrombosis was induced by laser injury of individual arterioles (WT: 52 injuries from 4 mice; *Lgals9^-/-^*: 62 injuries from 4 mice). Thrombus formation was monitored by confocal microscopy for 3 minutes per injury using an upright spinning-disk confocal microscope (Olympus) fitted with a 40× water-immersion objective (Nikon). Images were analyzed using Slidebook 6 (Intelligent Imaging Innovations) and ImageJ (National Institutes of Health), as previously described.^1,2^

#### Tail bleed assay

Mice (8–10 weeks old, all male) were anesthetized with intraperitoneal ketamine (76 mg/kg) and medetomidine (1 mg/kg) and placed prone on a heating pad. A 3 mm segment of the distal tail tip was excised, and the tail immediately immersed in 50 mL pre-warmed (37°C) saline for 20 min. Over this period, initial hemostasis (first cessation of flow lasting ≥10 s) and the timing of any re-bleeding episodes were recorded. The tube was then centrifuged at 1000 × *g* to pellet shed red blood cells; the supernatant was discarded and the pellet resuspended in 2 mL water to lyse erythrocytes. A hemoglobin standard curve was prepared in parallel by lysing a reference mouse blood sample (200, 100, 50, 20, 10, 5, and 1 µL) in 2 mL water. Samples and standards (100 µL each) were loaded onto a 96-well plate, and absorbance was read at 405 nm and interpolated against the standard curve to calculate blood loss.

#### Washed platelet preparation and activation

Whole blood was collected into acid-citrate-dextrose (ACD) anticoagulant (1:10) and centrifuged at 200 × *g* for 20 min to obtain platelet-rich plasma (PRP). PRP was treated with prostacyclin (PGI_2_) to prevent activation and centrifuged at 1000 × *g* for 10 min to pellet platelets. The resulting platelet pellet was resuspended in Tyrode buffer supplemented with ACD and prostacyclin, centrifuged at 1000 × *g*, and resuspended again in Tyrode buffer. Platelet counts were determined using a Sysmex analyzer and adjusted to 2 × 10^8^ platelets/mL.

Washed platelets (2 × 10^8^ cells/mL) were stimulated with vehicle (Ca^2+^/Mg^2+^-free PBS), recombinant Gal-9 (100, 300, or 600 nM; R&D Systems, Bio-Techne), or cross-linked collagen-related peptide (CRP-XL, 10 µg/mL; CambCol Laboratories) for 30 min at 37°C. Platelet activation was assessed by flow cytometry using FITC-conjugated PAC-1 (CD41/CD61; Invitrogen), PE-conjugated anti-P-selectin (BioLegend), and PE-Cy7-conjugated Annexin V (BioLegend), followed by fixation in 2% formalin (CellPath Ltd.). Samples were acquired on a CytoFlex S flow cytometer (Beckman Coulter) and analyzed using FlowJo (v10.8.1).

Intracellular calcium mobilization was assessed by flow cytometry. Washed platelets (2 × 10^8^ cells/mL) were loaded with Fluo-4 AM (cell permeant; F14217, Invitrogen) at 1 µM for 30 min at 37°C in the dark and treated with increasing concentrations of Gal-9 (100, 300, or 600 nM) for 10 min. Fluo-4 fluorescence was then acquired on a CytoFlex S flow cytometer (Beckman Coulter) for 2 min per sample and analyzed using FlowJo (v10.8.1); Fluo-4 median fluorescence intensity (MFI) was used as an index of intracellular Ca^2+^ influx.

Platelet aggregation was measured using a PAP-8E aggregometer (Bio/Data Corporation) under stirring conditions. Platelets were pre-warmed for 5 min, stirred for 2 min, and aggregation initiated with thrombin (0.01 U/mL; Merck Sigma-Aldrich). After 5 min, increasing concentrations of Gal-9 (100, 300, or 600 nM) were added and aggregation monitored for a further 5 min. Aggregation traces, maximal amplitude (Amax), and area under the curve (AUC) were calculated using PAP-8E software.

#### Endothelial cell activation

Human umbilical vein endothelial cells (HUVECs) were freshly isolated and cultured in M199 medium (Gibco) supplemented with 20% fetal bovine serum (FBS) and epidermal growth factor (EGF) (10 ng/mL) and used between passages P2–P4. Experiments were performed using HUVECs from four independent umbilical cord donors, with each donor representing a biological replicate.

For flow cytometry, cells were seeded in 60 mm petri dishes; for immunofluorescence, cells were seeded onto 0.2%-gelatin-coated ibidi µ-Slide VI 0.4 channels. Upon reaching ~90–95% confluence, cells were stimulated with TNF-α (tumor necrosis factor; 100 U/mL; R&D Systems) and IFN-γ (interferon gamma; 20 ng/mL; R&D Systems) for 24 h at 37°C, 5% CO_2_.

For flow cytometry, cells were gently scraped, washed with PBS, fixed in 2% formalin for 15 min, and permeabilized for 10 min (Permeabilization Buffer, FC005; R&D Systems). ICAM-1 was labeled on live cells prior to fixation using an APC-conjugated anti-ICAM-1 antibody (559771, BD Biosciences). Following blocking with 4% bovine serum albumin (BSA) in PBS for 1 h at room temperature, cells were incubated with primary anti-Gal-9 antibody (AF2045, R&D Systems; 1:75) and, after washing, AF488-conjugated donkey anti-goat secondary antibody (1:200; Invitrogen) for 90 min. Data were acquired on a CytoFlex S flow cytometer (Beckman Coulter) and analyzed using FlowJo (v10.8.1).

For immunofluorescence, cells in ibidi µ-Slide VI 0.4 channels were fixed and permeabilized as above, blocked, and stained with the same primary and secondary Gal-9 antibodies at the same dilutions. Images were acquired on an EVOS FL2 Auto (Invitrogen) microscope using a 20× objective.

#### Rotational thromboelastometry

Citrated whole blood was analyzed on a ROTEM instrument (TEM Innovations GmbH/Instrumentation Laboratory) at 37°C for a maximum of 60 minutes, assessing both extrinsic (EXTEM: recombinant tissue factor/phospholipid) and intrinsic (INTEM: ellagic acid/phospholipid contact activator) pathways, each combined with star-tem recalcification reagent. Samples were recalcified and spiked with recombinant Gal-9 (100 nM) or vehicle immediately before loading into ROTEM cups, and analysis was started without delay. Clotting time (CT, s) and maximum clot firmness (MCF, mm) were analyzed for each channel.

#### Human neutrophil isolation and NETosis assay

Human neutrophils were isolated from whole blood collected from healthy volunteers into EDTA tubes and processed within 30 min. Whole blood was layered onto a discontinuous Histopaque 1119/1077 (Merck Sigma-Aldrich) gradient and centrifuged at 400 × *g* for 25 min at room temperature with zero acceleration/brake. The neutrophil layer was carefully collected, topped up with ice-cold PBS-BSA buffer (Ca^2+^/Mg^2+^-free PBS, 0.5% BSA, 2 mM EDTA), and centrifuged at 350 × *g* for 10 min at 4°C. The pellet was resuspended in PBS, counted, and adjusted to 6.66 × 10^5^ cells/mL; all steps following the first centrifugation were performed at 4°C to minimize basal neutrophil activation and NETosis.

Human neutrophils were seeded onto poly-L-lysine-coated (0.1 mg/mL) 24-well plates and incubated for 30 min at 37°C, 5% CO_2_. Untreated or Gal-9-treated platelets (2 × 10^8^ cells/mL, final concentration) were then added and co-incubated with neutrophils for 3 h at 37°C. Cells were fixed in 2% formalin, washed with PBS, blocked with 2% BSA in PBS for 30 min, and permeabilized for 10 min. Cells were incubated for 1 h at room temperature with primary antibodies against myeloperoxidase (MPO, 3 µg/mL; MCA1757, Bio-Rad) and citrullinated histone H3 (CitH3; Abcam ab5103, 3 µg/mL), followed by AF488-conjugated anti-mouse IgG (MPO) and AF647-conjugated anti-rabbit IgG (CitH3), each at 1:500 for 1 h at room temperature. Following secondary antibody incubation, nuclei were counterstained with Hoechst dye (Thermo Fisher Scientific, #33342) for 15 min. After washing three times with 0.1% TBS-Tween 20 (3 min each), images were acquired on an EVOS FL microscope at 20× magnification. Images were analyzed in Fiji/ImageJ: MPO and CitH3 channels were individually thresholded, and the area of overlap between MPO-positive and CitH3-positive pixels (MPO+CitH3) was calculated and expressed as a percentage of total imaged area to quantify NET area.

#### Whole blood perfusion microfluidic assay

Glass coverslips were coated with recombinant Gal-9 (600 nM), collagen (50 µg/mL), von Willebrand factor (vWF, 100 µg/mL), or combinations of Gal-9 with collagen or vWF, for 90 min at room temperature, generating protein microspots (~3 mm diameter). Microspots were subsequently blocked with HEPES-BSA buffer for a further 90 min. Whole blood was recalcified (7.5 mM CaCl_2_, 3.75 mM MgCl_2_) and incubated with fibrinogen-AF488 (F13191, Invitrogen; 20 µg/mL), P-selectin-PE (BioLegend; 1:100), and Annexin V-AF647 (BioLegend; 1:100) for 10 min prior to perfusion. Blood was perfused over microspots in a Maastricht flow chamber at 100 s^-1^ or 1000 s^-1^, as previously described.^3^ In some experiments Gal-9 was blocked with 200 µg/mL of 1g3 clone of anti-Gal-9 antibody (Absolute Antibody) or isotype control antibody (Absolute Antibody). In some experiments, recombinant Gal-9 was added directly to recalcified whole blood, which was then incubated with the same antibody/fibrinogen panel for 30 min at 37°C prior to perfusion over collagen-coated (50 µg/mL) slides. Real-time multicolor fluorescence images were acquired every 3 minutes over 15 minutes (venous shear) or every 1 minute over 5 minutes (arterial shear) using an EVOS FL microscope, and platelet and fibrin coverage were quantified from all images using Fiji/ImageJ.

For perfusion over endothelial cells, HUVECs were seeded onto 0.2%-gelatin-coated ibidi µ-Slide VI 0.4 channels and, upon reaching near confluence, stimulated with TNF-α (100 U/mL) and IFN-γ (20 ng/mL) for 24 h. The endothelial monolayer was blocked with 2% BSA in M199 medium (without any supplements) for 1 h. In some experiments stimulated endothelial cells were blocked with 200 µg/mL of 1g3 clone of anti-Gal-9 antibody or isotype control antibody with 2% BSA in M199 medium for 1 hour. Recalcified whole blood (6.6 mM CaCl_2_, 2.1 mM MgCl_2_) was incubated with CD41-FITC (5B12, Agilent Technologies) (1:100) and fibrinogen-AF647 (F35200, Invitrogen) (20 µg/mL) for 10 min before perfusion over the endothelial monolayer for 5 min at a venous shear rate of 100 s^-1^. Real-time multicolor fluorescent images were acquired every 15 seconds over 5 min using an EVOS FL2 Auto microscope, and platelet and fibrin coverage were quantified from all images using Fiji/ImageJ.

#### Immunofluorescence of murine thrombi

Murine thrombi were harvested and immediately embedded in cryomold for snap-freezing on dry ice. Cryosections (5 µm) were cut, fixed in acetone, and blocked in 10% FBS/PBS. Two separate staining panels were performed with anti-mouse antibodies. For thrombus composition, sections were stained with directly conjugated antibodies against CD41 (PE, clone MWReg30, BioLegend; 1:100) and fibrinogen (FITC, F0111, Agilent Technologies; 1:100), alongside unconjugated primary antibody against Gal-9 (MAB3535, R&D Systems; 2 µg/mL) followed by an AF647-conjugated anti-rat secondary antibody. Isotype-matched controls were used for Gal-9 (rat IgG isotype control, MAB0061, R&D Systems; matched concentration) and fibrinogen (FITC-conjugated rabbit IgG isotype control, NBP2-24892, Novus Biologicals; matched concentration). For NETosis assessment, a separate set of sections was stained with primary antibodies against MPO (AF3667, R&D Systems; 3 µg/mL) and CitH3 (ab5103, Abcam; 3 µg/mL), followed by AF488-conjugated anti-goat and AF647-conjugated anti-rabbit secondary antibodies, respectively. Nuclei were visualized by Hoechst counterstain in both panels. Slides were imaged on an Invitrogen EVOS S1000 Spatial Imaging System at 20× magnification.

Image analysis was performed in Fiji/ImageJ. Thrombus area was defined by manual ROI tracing, and CD41-, fibrinogen-, and Gal-9-positive areas were quantified by automated thresholding as a percentage of total thrombus area. NETosis was quantified as the MPO^+^CitH3^+^ co-positive area, generated by combining individually thresholded MPO and CitH3 binary masks (Image Calculator AND function) and expressed as a percentage of total thrombus area. ≥3 non-adjacent sections were analyzed per thrombus, with data reported as mean ± SEM per animal.

### Supplemental Tables

**Table S1. Clinical characteristics of DVT patients**

| **Characteristic** | **Baseline (n=60)** | **Follow-up (n=59)** |
| --- | --- | --- |
| Age, years | 55.0 (32.8–59.3) [48.6 ± 14.6] | - |
| Age range, years | 22.4–76.2 | - |
| Male sex — n (%) | 34 (56.7%) | - |
| Female sex — n (%) | 26 (43.3%) | - |
| **DVT Location** |  |  |
| Distal (infrapopliteal) — n (%) | 26 (43.3%) | - |
| Popliteal — n (%) | 12 (20.0%) | - |
| Femoral — n (%) | 13 (21.7%) | - |
| Iliac/pelvic — n (%) | 7 (11.7%) | - |
| Caval — n (%) | 2 (3.3%) | - |
| **Laboratory Values** |  |  |
| Hemoglobin, g/dL | 14.0 (12.4–14.9) [13.6 ± 1.8] | 14.6 (13.7–15.4) [14.2 ± 1.7] |
| CRP, mg/dL | 0.85 (0.22–3.62) [2.56 ± 4.81] | 0.14 (0.09–0.28) [0.24 ± 0.30] |
| Creatinine, mg/dL | 0.85 (0.75–0.97) [0.89 ± 0.24] | 0.86 (0.75–0.99) [0.88 ± 0.19] |
| Platelet count, ×10^9^/L | 277.5 (217.5–327.0) [287.8 ± 99.3] | 248.0 (210.5–299.0) [247.7 ± 63.8] |

*Values are median (IQR) [mean ± SD] or n (%). CRP, C-reactive protein; IQR, interquartile range; SD, standard deviation.*

### Major Resources Table

**Major Resources Table. Key reagents, antibodies, animals and primary human cells,** grouped by the assay in which they were used. Working concentrations and in vivo doses are given in the Supplemental Methods.

#### A. Antibodies used for flow cytometry

| **Target** | **Conjugate** | **Vendor** | **Catalogue / clone** | **Species raised in** |
| --- | --- | --- | --- | --- |
| CD41/CD61 (PAC-1) | FITC | Invitrogen | PAC-1 | Mouse |
| P-selectin | PE | BioLegend | 304906 (clone AK4) | Mouse |
| Annexin V | PE-Cy7 | BioLegend | Not stated | Not applicable (recombinant binding protein) |
| ICAM-1 | APC | BD Biosciences | 559771 (clone HA58) | Mouse |
| Galectin-9 | Unconjugated (primary) | R&D Systems | AF2045 | Goat |
| Donkey anti-goat IgG (secondary, Gal-9) | AF488 | Invitrogen | A11055 | Donkey |

#### B. Antibodies used for immunofluorescence and intravital imaging

| **Target** | **Conjugate** | **Vendor** | **Catalogue / clone** | **Species raised in** |
| --- | --- | --- | --- | --- |
| GPIbβ/CD42c (intravital, IVC) | DyLight 649 | Emfret Analytics | X-649 | Rat |
| Ly6G (intravital, IVC) | PE | BioLegend | 1A8 | Rat |
| GPIbβ (laser arterial injury) | DyLight 649 | Emfret Analytics | X-649 | Rat |
| Galectin-9 (HUVEC IF) | Unconjugated (primary) | R&D Systems | AF2045 | Goat |
| Donkey anti-goat IgG (secondary, HUVEC IF) | AF488 | Invitrogen | A11055 | Donkey |
| Myeloperoxidase (neutrophil–platelet co-culture) | Unconjugated (primary) | Bio-Rad | MCA1757 | Mouse |
| Citrullinated histone H3 (neutrophil–platelet co-culture) | Unconjugated (primary) | Abcam | ab5103 | Rabbit |
| Anti-mouse IgG (secondary, MPO) | AF488 | Invitrogen | A11029 | Goat |
| Anti-rabbit IgG (secondary, CitH3) | AF647 | Invitrogen | A21245 | Goat |
| CD41 (murine thrombus composition) | PE | BioLegend | MWReg30 | Rat |
| Fibrinogen (murine thrombus composition) | FITC | Agilent Technologies | F0111 | Rabbit |
| Galectin-9 (murine thrombus composition) | Unconjugated (primary) | R&D Systems | MAB3535 | Rat |
| Anti-rat IgG (secondary, Gal-9) | AF647 | Invitrogen | A48272 | Donkey |
| Rat IgG isotype control (matched to MAB3535) | Unconjugated | R&D Systems | MAB0061 | Rat |
| Rabbit IgG isotype control (matched to F0111) | FITC | Novus Biologicals | NBP2-24892 | Rabbit |
| Myeloperoxidase (murine thrombus NETosis panel) | Unconjugated (primary) | R&D Systems | AF3667 | Goat |
| Citrullinated histone H3 (murine thrombus NETosis panel) | Unconjugated (primary) | Abcam | ab5103 | Rabbit |
| Anti-goat IgG (secondary, MPO) | AF488 | Invitrogen | A11055 | Donkey |
| Anti-rabbit IgG (secondary, CitH3) | AF647 | Invitrogen | A21245 | Goat |

#### C. Antibodies used in whole blood perfusion (flow chamber) assays

| **Target** | **Conjugate** | **Vendor** | **Catalogue / clone** | **Species raised in** |
| --- | --- | --- | --- | --- |
| Fibrinogen (Maastricht flow chamber) | AF488 | Invitrogen | F13191 | Not applicable (labelled human fibrinogen) |
| P-selectin (Maastricht flow chamber) | PE | BioLegend | 304906 (clone AK4) | Mouse |
| Annexin V (Maastricht flow chamber) | AF647 | BioLegend | Not stated | Not applicable (recombinant binding protein) |
| Galectin-9 (blocking, flow chamber) | Unconjugated | Absolute Antibody | Ab04373-10.0 (clone 1G3) | Human IgG1 (recombinant; parent hybridoma mouse) |
| Isotype control (matched to 1g3) | Unconjugated | Absolute Antibody | Ab00178-10.0 | Human IgG1 |
| CD41 (HUVEC perfusion assay) | FITC | Agilent Technologies | 5B12 | Mouse |
| Fibrinogen (HUVEC perfusion assay) | AF647 | Invitrogen | F35200 | Not applicable (labelled human fibrinogen) |
| Galectin-9 (blocking, HUVEC perfusion) | Unconjugated | Absolute Antibody | Ab04373-10.0 (clone 1G3) | Human IgG1 (recombinant; parent hybridoma mouse) |
| Isotype control (matched to 1g3, HUVEC perfusion) | Unconjugated | Absolute Antibody | Ab00178-10.0 | Human IgG1 |

#### D. Other key immunodetection and functional reagents

| **Reagent** | **Vendor** | **Catalogue** | **Used in** |
| --- | --- | --- | --- |
| Recombinant human galectin-9 | R&D Systems, Bio-Techne | 2045-GA | Platelet stimulation (100–600 nM), flow chamber coating, ROTEM spiking |
| Cross-linked collagen-related peptide (CRP-XL) | CambCol Laboratories | Not stated | Positive control for platelet activation (10 µg/mL) |
| Fluo-4 AM (cell permeant) | Invitrogen | F14217 | Platelet intracellular calcium mobilization (1 µM) |
| Hoechst 33342 | Thermo Fisher Scientific | 33342 | Nuclear counterstain (NETosis and thrombus panels) |
| Permeabilization buffer | R&D Systems | FC005 | HUVEC flow cytometry and immunofluorescence |
| Histopaque 1119 / 1077 | Merck Sigma-Aldrich | 11191 (Histopaque-1119) / 10771 (Histopaque-1077) | Human neutrophil isolation |
| EXTEM / INTEM / star-tem reagents | TEM Innovations / Instrumentation Laboratory | 503-05 (ex-tem), 503-02 (in-tem), 503-01 (star-tem) | Thromboelastometry (ROTEM) |
| Human Luminex Discovery Assay (magnetic bead-based multiplex immunoassay, galectin-9 analyte) | R&D Systems, Bio-Techne | LXSAHM | Plasma galectin-9 quantification in patients and controls (Figure 1) |
| Recombinant human TNF-α | R&D Systems, Bio-Techne | 210-TA | HUVEC stimulation (100 U/mL, 24 h) |
| Recombinant human IFN-γ | R&D Systems, Bio-Techne | 285-IF | HUVEC stimulation (20 ng/mL, 24 h) |

#### E. Animals and primary human cells

| **Model** | **Strain / designation** | **Source** | **Persistent identifier** | **Used in** |
| --- | --- | --- | --- | --- |
| Wild-type mouse | C57BL/6J | The Jackson Laboratory | RRID:IMSR_JAX:000664 (JAX stock #000664) | IVC stenosis DVT model (male and female); intravital microscopy and tail bleeding assay (male only); 8–10 weeks |
| Gal-9-deficient mouse | B6(FVB)-*Lgals9^tm1.1Cfg^*/Mmucd | Mutant Mouse Resource and Research Center (MMRRC), University of California, Davis | RRID:MMRRC_031952-UCD | IVC stenosis DVT model (male and female); intravital microscopy and tail bleeding assay (male only); 8–10 weeks |
| Primary human endothelial cells | HUVEC, four independent umbilical cord donors, passages P2–P4 | Freshly isolated (REC 15/NE/0285) | Not applicable | Endothelial activation, immunofluorescence, whole blood perfusion |
| Primary human neutrophils and platelets | Healthy volunteer donors | University of Birmingham (ERN_11-0175) | Not applicable | Platelet activation, NETosis, flow chamber assays |

*The Gal-9 neutralizing antibody (clone 1g3) and its matched isotype control were provided by Absolute Antibody. D-dimer was measured by a quantitative latex agglutination assay (central laboratory, Medical University of Vienna; LLOQ 0.27 mg/L).*

### Supplemental Figures

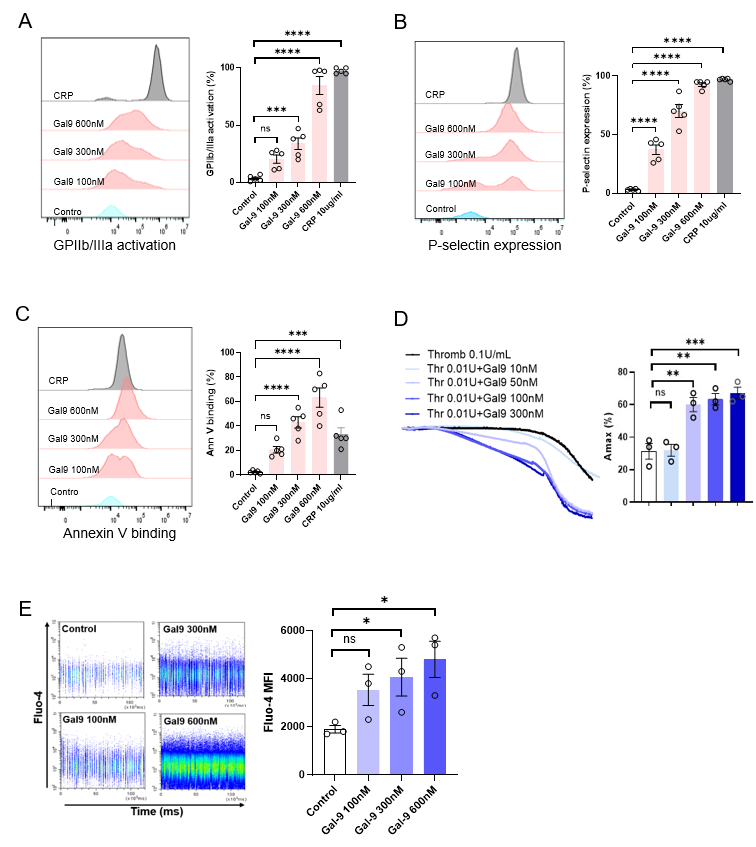

**Figure S1. Gal-9 directly modulates platelet activation**

(A–C) Washed platelets were stimulated with vehicle, recombinant Gal-9 (100, 300 or 600 nM) or CRP-XL (10 µg/mL) and analyzed by flow cytometry. Gal-9 dose-dependently increased (A) GPIIb/IIIa activation (PAC-1 binding), (B) P-selectin surface expression and (C) phosphatidylserine exposure (annexin V binding) (n = 5). (D) Gal-9 potentiated aggregation of washed platelets triggered by thrombin (0.01 U/mL) (n = 3). (E) Fluo-4-loaded human washed platelets were treated with increasing concentrations of Gal-9 for 10 min and fluorescence was acquired by flow cytometry for 2 min; increased Fluo-4 MFI indicates increased intracellular Ca^2+^ influx (n = 3). Data are mean ± standard error of the mean (SEM) from n independent donors; one-way ANOVA with Tukey’s multiple comparison test. **P* < .05, ***P* < .01, *****P* < .0001; ns, not significant.

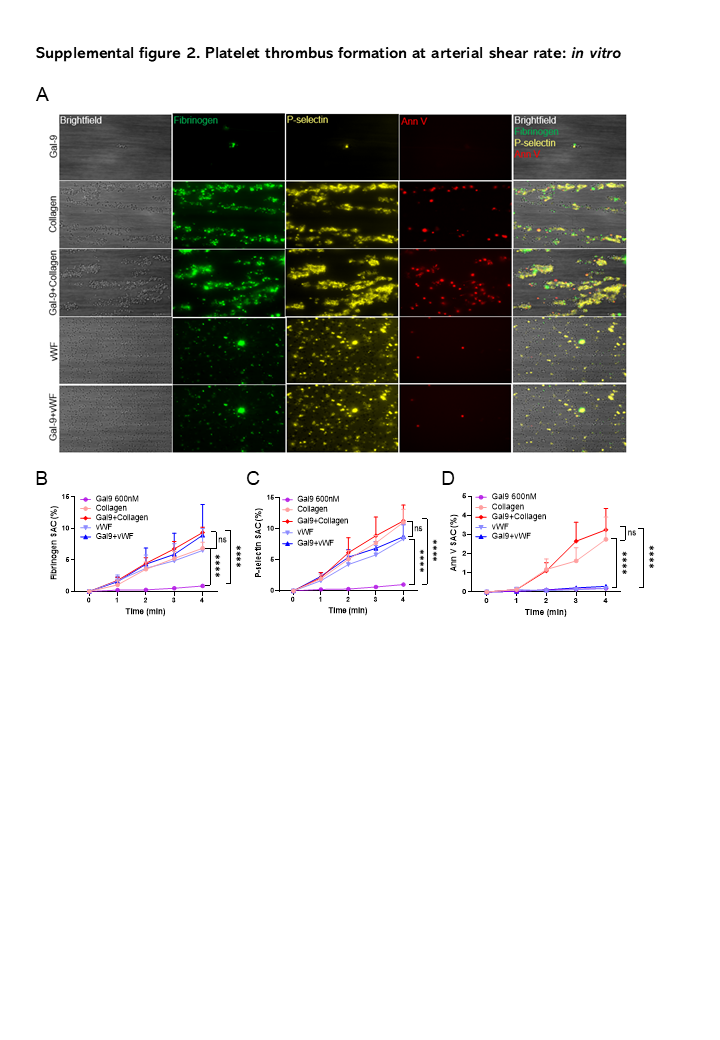

**Figure S2. Platelet thrombus formation at arterial shear rate *in vitro***

(A) Representative multichannel images at 5 minutes of recalcified human whole blood perfused at arterial shear rate (1000 s^-1^) over microspots coated with immobilized Gal-9 (600 nM), collagen (50 µg/mL), von Willebrand factor (vWF, 100 µg/mL), or combinations thereof (Gal-9 + collagen; Gal-9 + vWF). (B–D) Platelet activation over 5 minutes, expressed as surface area coverage (SAC, %): (B) fibrinogen binding (AF488), (C) P-selectin exposure (PE), and (D) phosphatidylserine exposure (annexin V-AF647) (n = 4 biological replicates). Scale bar = 50 µm. Data are mean ± SEM; two-way ANOVA with Tukey’s multiple comparison test. **P* < .05, ***P* < .01, *****P* < .0001; ns, not significant.

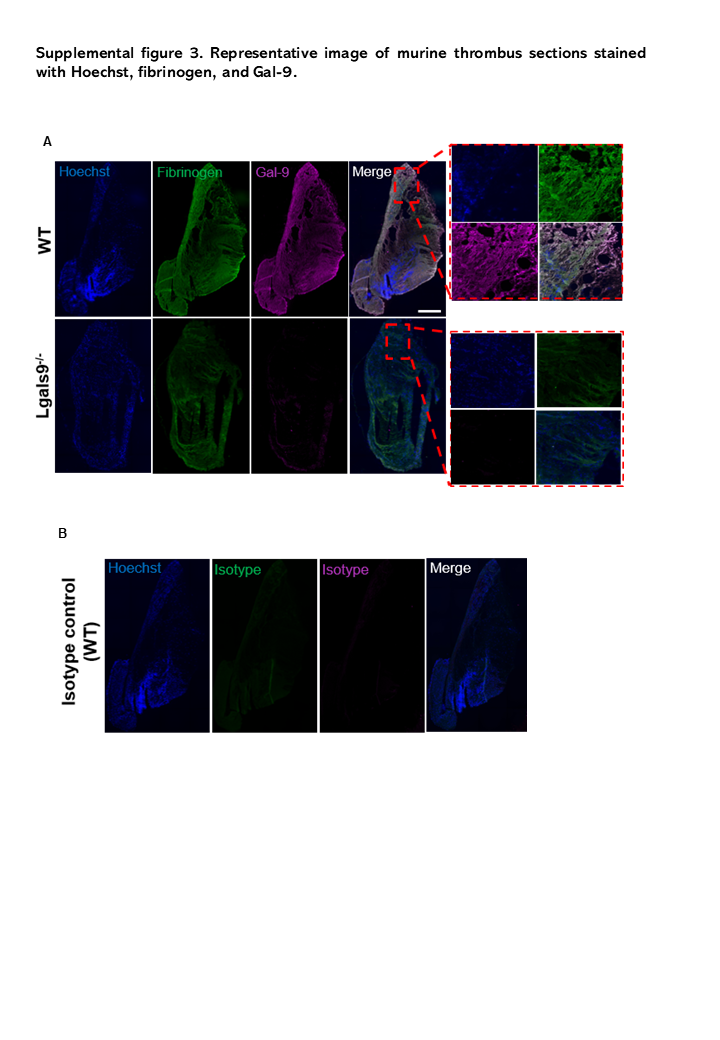

**Figure S3. Co-immunostaining of fibrinogen and** **Gal-9 in murine thrombus**

Immunofluorescence images of thrombus cryosections. (A) Representative images of thrombus co-stained for Hoechst (blue), fibrinogen (green), and Gal-9 (magenta), with merged channels, in WT (top) and *Lgals9^-/-^* (bottom) mice, showing Gal-9 localization within the platelet-rich thrombus; red dashed boxes indicate the regions shown at higher magnification on the right. (B) Isotype control of the same WT thrombus. Scale bar = 500 µm.

### Supplemental Video Legends

**Video S1. IVC stenosis model of DVT**

Representative corresponding intravital video (Figure 2H) showing platelet and neutrophil recruitment to the vessel wall following 3 hours of IVC stenosis.

**Video S2. Laser-injury arterial thrombosis**

Representative corresponding video (Figure 2K) showing platelet thrombus formation following laser-induced arterial injury.
